# The AUUUC repeat RNA aggregates sequester RNA-binding proteins like NOVA2 and lead to iron dyshomeostasis in spinocerebellar ataxia type 37

**DOI:** 10.1101/2025.09.03.673996

**Authors:** Ana S. Figueiredo, Joana R. Loureiro, Paula Sampaio, Hugo Osório, Montserrat Soler-López, José Bessa, Sylvain Bohic, Sandra Macedo-Ribeiro, Isabel Silveira

## Abstract

Transcribed nucleotide repeat expansions can contribute to disease by altering RNA structure and function. Spinocerebellar ataxia type 37 (SCA37) is a neurodegenerative disorder caused by a pathogenic ATTTC repeat insertion within a non-pathogenic ATTTT repeat in the 5’ untranslated region of *DAB1*. We have shown that the AUUUC repeat RNA forms aberrant nuclear aggregates in cells, subsequently confirmed by others in neurons from subjects with familial adult myoclonic epilepsy carrying a similar ATTTC repeat insertion. However, the mechanism by which these RNA aggregates cause neuropathology remains unknown. Here, we show that overexpression of the ATTTC repeat in human neural stem cells leads to the formation of abnormal nuclear RNA aggregates, supporting an AUUUC repeat-mediated mechanism of pathology through the sequestration of RNA-binding proteins (RBP). We identified 12 AUUUC repeat-interacting RBPs with specific neuronal functions, including NOVA2, which we demonstrate to colocalize with the AUUUC repeat aggregates. Moreover, we further investigated the accumulation of iron in these aggregates and observed a significant colocalization of iron and NOVA2 hotspots in ATTTC repeat-expressing cells, a pattern absent in control cells. Together, these findings uncover a novel RNA-mediated mechanism of pathology involving both RBPs and iron, expanding the current understanding of RNA repeat toxicity.

## INTRODUCTION

Short tandem repeat (STR) sequences are polymorphic regions of the DNA composed of tandem repeat units of 1-6 bp, which constitute about 7% of the human genome (1). Expanded STRs with repeat length above the pathogenicity threshold are known to be linked to over 60 developmental, neurodegenerative, or neuromuscular diseases, including Huntington’s disease (HD), myotonic dystrophy types 1 and 2 (DM1 and DM2), frontotemporal dementia/amyotrophic lateral sclerosis (FTD/ALS) and several spinocerebellar ataxias (SCAs) (1,2). Together, these data emphasize the crucial role of DNA/RNA STR sequence length, as well as repeat motifs, in cell biology and disease.

SCAs represent a diverse group of disorders, both clinically and genetically, typically marked by adult-onset progressive ataxia affecting gait, limb coordination, and speech due to cerebellar neuronal degeneration, often with associated peripheral neuropathy (3,4). The prevalence of autosomal-dominant SCAs differs significantly across regions worldwide, ranging from 1.6 to 5.6 cases per 100,000 individuals (5). While many SCAs result from CAG repeat expansions in exonic gene regions encoding polyglutamine tracts, some are caused by expansions of other types of STRs in non-coding gene sequences, like the pentanucleotide repeat insertions causing SCA10, SCA31 and SCA37 (6–9).

Spinocerebellar ataxia type 37 (SCA37) is a progressive neurodegenerative disorder characterized by initial signs of gait ataxia, dysarthria, and abnormal ocular movements (7,10–12). These symptoms progress to a very debilitating condition for which no curative treatments are currently available.

Our previous studies identified a pathogenic ATTTC repeat insertion within an ATTTT repeat sequence in a 5′UTR intron of the *DAB1* gene as the genetic cause of SCA37 (7). *DAB1* encodes the reelin adaptor protein, which plays a crucial role in neurodevelopmental processes, such as neuronal migration, brain layer organization, and axonal pathfinding (13–17). In unaffected individuals, the *DAB1* gene contains an (ATTTT)_n_ sequence ranging from 7 to 400 ATTTT units, while affected individuals carry the pathogenic insertion with the configuration [(ATTTT)_60-79_(ATTTC)_31-100_(ATTTT)_58-415_] (7,10,12). Notably, the length of the (ATTTC)_n_ insertion is inversely correlated with the age of disease onset, and this pentanucleotide repeat shows length instability, particularly when transmitted paternally (7). Since our finding of this (ATTTC)_n_ insertion in *DAB1* underlying SCA37, similar repeat insertions in seven genes have been linked to neurological disorders classified as familial adult myoclonic epilepsies (FAME 1–4, 6, 7 and 8), affecting numerous families worldwide (18–24). Furthermore, this (ATTTC)_n_ tract was found to form dimeric DNA structures promoted by divalent ions (25). These findings underscore the pathogenic potential of the (ATTTC)_n_ insertions, which, when present in diverse genomic contexts, can lead to a range of diseases, suggesting a shared underlying mechanism.

In many neurodegenerative and neuromuscular diseases, these RNAs with expanded STR sequences are known to form abnormal nuclear foci, as they sequester RNA-binding proteins (RBPs) and disrupt normal cellular functions (26–30). Evidence to support this has been observed in 1) DM1, in which the transcription of (CUG)_n_ results in the sequestration of RBPs such as CELF and MBNL in autopsy brain tissue of DM1 patients, leading to significant downstream effects on alternative splicing and polyadenylation (30); 2) in fragile X-associated tremor/ataxia syndrome (FXTAS), expanded CGG RNA repeats sequester proteins like DROSHA and DGCR8 on brain tissue (hippocampal area) of FXTAS patients, thereby impairing microRNA processing (29); and 3) in ALS/FTD, transcription of *C9orf72* hexanucleotide repeats leads to the sequestration of ribonucleoproteins such as nucleolin in motor cortex tissue from patients carrying the *C9orf72* hexanucleotide expansion, triggering nucleolar stress and contributing to disease pathology (31). On the other hand, the huntingtin protein (HTT), which carries an expanded polyglutamine tract in Huntington’s disease-affected cells, is itself an RBP that interacts with several RNAs, like the long non-coding RNA *NET1*, influencing its cellular levels and paraspeckle formation (32). It is remarkable that most transcripts with which HTT interacts function in mitochondrial respiration, for which iron homeostasis is fundamental. Indeed, Chen et al., found that in the R6/2 Huntington’s disease mouse model, redox-active ferrous iron accumulates in perinuclear endocytic/lysosomal compartments of neurons alongside dysregulated iron homeostasis (33).

It is known that iron plays a critical role in the function of several RBPs, by acting as a structural cofactor (34). In RNA-binding domains, iron can stabilize protein structure or facilitate protein-RNA interactions, not only influencing the folding and stability of RBPs but also acting as molecular switches that modulate RNA affinity (34,35). Importantly, iron dyshomeostasis has been linked to several neurodegenerative disorders, including Alzheimer’s disease, Parkinson’s disease, and multiple sclerosis (36–38). Both iron accumulation and deficiency can contribute to neuronal death through mechanisms such as oxidative stress, membrane disruption, and mitochondrial dysfunction (38).

We have previously demonstrated that SCA37 AUUUC repeat RNA forms aberrant nuclear aggregates in non-neural human cells and induces lethal malformations and developmental defects in zebrafish (7). These AUUUC repeat RNA aggregates have been confirmed in cerebellar Purkinje cells and cortical neurons from FAME-affected subjects (22). Altogether, these data suggest that RNA toxicity is a fundamental mechanism for pathogenesis. However, how these RNA aggregates cause neuropathology remains unknown.

In this study, we demonstrate that the pathogenic AUUUC repeat RNA forms nuclear aggregates in human neural stem cells and exhibits increased binding affinity for RBPs with specific neuronal functions such as NOVA2, MATRIN-3 and PTBP1/3, compared with the non-pathogenic AUUUU repeat tract. Colocalization studies confirmed that NOVA2 colocalizes with the AUUUC repeat RNA in neural stem cells, reinforcing a model in which this pathogenic RNA sequesters RBPs, compromising their function. To support this mechanism, we show that these nuclear AUUUC repeat RNA-NOVA2 aggregates colocalize with iron hotspots in cells expressing the pathogenic ATTTC repeat, a pattern absent in control cells. This suggests that the AUUUC repeat RNA-protein complexes interfere with cellular iron homeostasis, potentially underlying SCA37 pathology.

## MATERIAL AND METHODS

### Cell culture

Human embryonic kidney 293T (HEK293T) cells (ATCC #CRL-3216) were cultured in DMEM GlutaMAX™ medium (#31966047, Gibco, ThermoFisher Scientific) with 10% fetal bovine serum (#A5256801, Gibco, ThermoFisher Scientific), 100 units/mL penicillin, and 0.1 mg/mL streptomycin (#15140122, ThermoFisher Scientific), at 37°C, with 5% CO_2_.

Human neural stem cells (hNSC; line CB192) (39) were maintained in DMEM-F12 GlutaMAX™ (#31331028, Gibco, ThermoFisher Scientific) supplemented with 1× N-2 (#17502048; Gibco, ThermoFisher Scientific,), 0.05× B27 (#17504044, Gibco, Thermofisher Scientific), 10 ng/ml EGF (#315-09, Peprotech); 10ng/ml FGF (#100-18B, Peprotech); 1 μg/ml Laminin (#L2020-1MG, Merck Life Sciences), and 100 units/mL penicillin + 0.1 mg/mL streptomycin (#15140122, Sigma Aldrich) in T-flasks or plates pre-coated with 0.01% Poly-L-Lysin (#P8920-100ML, Merck Life Sciences). The cells were maintained at 37°C, with 5% CO_2._

### RNA fluorescent *in situ* hybridization (RNA-FISH)

Human neural stem cells (hNSCs) were transfected with the pCDH-CMV-EF1-GFP-Puro vector (7), expressing the non-pathogenic N(ATTTT)_7_, N(ATTTT)_139_ or the pathogenic Ins(ATTTC)_58_ sequences in the configuration of (ATTTT)_57_(ATTTC)_58_(ATTTT)_71_, using the Lipofectamine 2000 (#11668027, Invitrogen) according to the manufacturer procedure. Forty-eight hours after transfection, the cells were fixed in 4% paraformaldehyde. Permeabilization was carried out in 2% acetone in PBS for 5 min at room temperature. Following this, the coverslips were incubated in pre-hybridization solution (30% formamide (#F9037-100ml, Sigma Aldrich), 2× SSC (#15557-044, Gibco, Thermofisher Scientific) for 10 min at room temperature, then subjected to hybridization in a solution containing 0.5 ng/μL Texas Red-labelled (GAAAU)_5_ probe (Integrated DNA Technologies, IDT), 30% formamide, 2× SSC, 0.02% BSA (#AM2616, Ambion), 66 μg/ml yeast tRNA (#15401011, Invitrogen), and 2 mM ribonucleoside vanadyl complex (#R3380-5ML, Sigma-Aldrich) at 37°C for 3 h. After two washing steps with 2× SSC (10 min and 2 min, respectively), cells were counterstained with DAPI (1:4000) for 10 min at room temperature and the coverslips were mounted in Ibidi mounting media (#50001, ibidi). FISH signal was acquired in Leica DMI6000 inverted epifluorescence microscope, using a HCX PL APO CS 63×/1.30 GLYC 21°C objective (Leica Microsystems).

### RNA pull-down

To identify candidate proteins interacting with the AUUUC repeat RNA, we performed an RNA pull-down assay using (AUUUC)₁₀ and (AUUUU)₁₀ RNA oligonucleotides synthesized by Exiqon. Total protein extracts from the human neuroblastoma cell line were mixed with the lysis buffer supplemented with protease inhibitors (50 mM Tris-HCl pH 7.5, 1 mM EDTA, 10% glycerol, 50 nM NaF, 5 mM sodium pyrophosphate, 1% Triton X 100, 1 mM DTT, 0.1 mM PMSF, Na_3_VO_4_ in a 1:100 dilution and protease inhibitor mix). RNA labelling (biotinylation), protein-RNA binding, and subsequent protein elution were performed using the Pierce™ Magnetic RNA-Protein Pull-Down Kit (#20164, Thermofisher Scientific), following the manufacturer’s instructions. Proteins eluted from the RNA pull-down assay were identified through liquid chromatography-mass spectrometry (LC-MS) and analyzed using the Proteome Discoverer software (Thermofisher Scientific). Differential protein interactions between (AUUUC)₁₀ and (AUUUU)₁₀ were statistically analyzed using ANOVA and proteins with a higher abundance ratio in the (AUUUC)₁₀ RNA pull-down were selected as candidate proteins specifically interacting with (AUUUC)_n_. Gene ontology analysis was conducted using the Functional Annotation Tool available in the Database for Annotation, Visualization and Integrated Discovery (DAVID; https://david.ncifcrf.gov/) for the 28 RBPs that showed increased binding to the (AUUUC)_10_ repeat. Gene expression analysis was performed using RNA-seq data from the Genotype-Tissue Expression (GTEx) (https://gtexportal.org) project to assess the expression patterns of the top 12 identified RBPs across multiple human brain regions or across different tissues for *DAB1* and *NOVA2*. Expression levels, measured in transcripts per million (TPM), are visualized in a heatmap.

### Liquid-chromatography and mass-spectrometry (LC-MS) analysis

Each sample was processed for proteomic analysis following the solid-phase-enhanced sample-preparation (SP3) protocol and enzymatically digested with Trypsin/LysC as previously described (40). Protein identification and quantitation were performed by nanoLC-MS/MS. This equipment is composed of an Ultimate 3000 liquid chromatography system coupled to a Q-Exactive Hybrid Quadrupole-Orbitrap mass spectrometer (Thermo Scientific, Bremen, Germany). Each sample was loaded onto a trapping cartridge (Acclaim PepMap C18 100 Å, 5 mm × 300 μm i.d., 160454, Thermo Scientific, Bremen, Germany) in a mobile phase of 2% ACN, 0.1% FA at 10 μL/min. After 3 min loading, the trap column was switched in-line to a 50 cm × 75 μm inner diameter EASY-Spray column (ES803, PepMap RSLC, C18, 2 μm, Thermo Scientific, Bremen, Germany) at 300 nL/min. Separation was achieved by mixing A: 0.1% FA and B: 80% ACN, 0.1% FA with the following gradient: 80 min (2.5% B to 40% B), 20 min (40% B to 95% B), and 20 min (hold 99% B). Subsequently, the column was equilibrated with 2.5% B for 27 min. Data acquisition was controlled by Xcalibur 4.0 and Tune 2.8 software (Thermo Scientific, Bremen, Germany).

The mass spectrometer was operated in the data-dependent (dd) positive acquisition mode alternating between a full scan (m/z 300-1750) and subsequent HCD MS/MS of the 10 most intense peaks from a full scan (normalized collision energy of 27%). The ESI spray voltage was 1.9 kV. The global settings were as follows: use lock masses best (m/z 445.12003), lock mass injection Full MS and chrom. peak width (FWHM) of 15 s. The full scan settings were as follows: 70 k resolution (m/z 200), AGC target 3 × 10^6^, maximum injection time 120 ms; dd settings: minimum AGC target 2.20 × 10^3^, intensity threshold 2.0 × 10^4^, charge exclusion: unassigned, 1, >6, peptide match preferred, exclude isotopes on, and dynamic exclusion 30 s. The MS2 settings were as follows: microscans 1, resolution 35 k (m/z 200), AGC target 1 × 10^5^, maximum injection time 110 ms, isolation window 2.0 m/z, isolation offset 0.0 m/z, dynamic first mass, and spectrum data type profile.

The raw data was processed using the Proteome Discoverer 2.2.0.388 software (Thermo Scientific) and searched against the UniProt database for the Homo sapiens taxonomic selection (2017_09). The Sequest HT search engine was used to identify tryptic peptides. The ion mass tolerance was 10 ppm for precursor ions and 0.02 Da for fragment ions. Maximum allowed missing cleavage sites was set to two. Cysteine carbamidomethylation was defined as constant modification. Methionine oxidation and protein N-terminus acetylation were defined as variable modifications. Peptide confidence was set to high. The processing node Percolator was enabled with the following settings: maximum delta Cn 0.05; decoy database search target false discovery rate 1%, validation based on q-value. Protein label-free quantitation was performed with the Minora feature detector node at the processing step. Precursor ions quantification was performed at the consensus step with the following parameters: unique plus razor peptides were considered, precursor abundance based on area, and normalization based on total peptide amount. For hypothesis testing, protein ratio calculation was pairwise ratio-based, and a t-test (background-based) hypothesis test was performed. The mass spectrometry proteomics data have been deposited to the ProteomeXchange Consortium via the PRIDE (41) partner repository with the dataset identifier PXD066760.

### Cloning a His-tag encoding sequence at the NOVA2 cDNA expressing vector

To design a NOVA2 expression vector with a histidine tag (His-tag), a 14 His-tag was inserted into the C-terminal of the NOVA2 protein. The C-terminal was selected due to the presence of a poly-glycine sequence, which provides sufficient flexibility to prevent interference with other amino acids of the protein, thereby avoiding the folding of the C-terminal into the protein core. To create the NOVA2-His-tag expression vector, the NOVA2 Human Untagged Clone (#SC303210, Origene) vector was modified by a His-tag sequence of 14 histidines inserted, in frame, at the C-terminal of the NOVA2 by the *in vivo* assembly (IVA) cloning method (42). The amplification of the NOVA2 Human Untagged Clone was performed using KOD DNA polymerase (#71085-3, Sigma-Aldrich). The reaction used 1 ng of template DNA vector, 1× KOD Master Mix, 7.5 μM each of the following primers: 5’-CATCATCATCATCATCATTGAGGCCTGTGGTGTGTGCTC-3’ and 5’-ATGATGATGATGATGATGTCCCACTTTCTGGGGGTTTGAGG-3’, 1.25 μL of 100% DMSO and nuclease-free H_2_O up to 25 μL. The PCR protocol began with a 2-minute denaturation at 98°C, followed by 25 cycles of 20 s at 98°C, 10 s at 60°C, and 4.5 min at 70°C, and concluded with a final 10 min extension at 70°C. The PCR product was then digested with the Fast Digest DpnI enzyme (#FD1703, Thermofisher Scientific). The digested material was transformed into DH5α competent bacteria, and colonies were purified and sequenced by Sanger Sequence to validate the success of His-tag cloning sequence insertion.

### RNA-FISH followed by Immunofluorescence (IF)

hNSCs were seeded in glass coverslips in 24-well plates at a density of 120,000 cells/well. On following day, cells were co-transfected with the pCDH-CMV-EF1-GFP-Puro vector (7), expressing the non-pathogenic N(ATTTT)_7_, N(ATTTT)_139_ or the pathogenic Ins(ATTTC)_58_ sequences and the NOVA2-His-tag expression vector, using the Lipofectamine 2000 (#11668027, Invitrogen) according to the manufacturer procedure. Forty-eight hours after transfection, the cells were fixed in 4% paraformaldehyde. Permeabilization and FISH were performed as previously described in the section “RNA Fluorescent *in situ* hybridization (RNA-FISH)”. After the two washes with 2× SSC, coverslips were blocked with 2.5% BSA for 1 h at room temperature. Subsequently, the primary Anti-His-tag antibody (#05-949, Millipore) was diluted to 1:1500 in the blocking solution, and coverslips were incubated with the primary antibody overnight at 4°C. The next day, coverslips were washed three times with 1× PBS and incubated with the secondary antibody (Alexa 647 Rabbit Anti-Mouse, diluted at 1:1500 in 1× PBS) for 1 h at room temperature. Finally, coverslips were counterstained with DAPI (diluted 1:4000) for 10 min at room temperature and mounted using ibidi mounting media (#50001, ibidi). FISH and IF signals were visualized using a Leica Microsystems TCS SP5 II confocal microscope with a HC PL APO Lbl. Blue 63× /1.40 Oil objective (Leica Microsystems).

### Analysis of NOVA2 localization in human neural stem cells (hNSC)

To quantify the intensity of NOVA2, each cell nucleus was segmented into radial sections 10 pixels wide and the intensity of NOVA2 was quantified within each section. A total of 69 cells transfected with the non-pathogenic N(ATTTT)_7_, 61 cells transfected with the N(ATTTT)_139_ and 74 cells transfected with the pathogenic Ins(ATTTC)_58_ repeat were analyzed using an in-house macro developed to quantify NOVA2 intensity across cells, ensuring consistent and reproducible measurements (doi: 10.5281/zenodo.15790995). To account variations in cell size, we defined the “nuclear periphery” and “nuclear core” based on the diameter of each nucleus. Specifically, for cells with a diameter greater than 60 pixels, the “peripheral” region was defined as the inner 20 pixels from the nuclear membrane toward the center. Conversely, for cells with a diameter of less than 60 pixels, the “peripheral” region was defined as the inner 10 pixels from the nuclear membrane toward the center. The intensity of the signal in the periphery or core region of the nucleus between the different conditions was compared using a two-way ANOVA with multiple comparison test, with p<0.05 considered statistically significant. For comparing NOVA2 intensity between the periphery and core for each condition, a Mann-Whitney U test was performed with p<0.05 considered statistically significant.

### Cell seeding and transfection of HEK293T cells in Si_3_Ni_4_ membranes

HEK293T cells were plated in 24-well plates with a density of 35,000 cells/well, and co-transfected with pCDH-CMV-EF1-GFP-Puro vector (7), encoding either the non-pathogenic N(ATTTT)_7_ or the pathogenic Ins(ATTTC)_58_ sequences, and with the NOVA2-His-tag plasmid, using JetPrime® transfection reagent (#101000027, Polyplus), according to the manufacturer’s instructions. Mock-transfected cells underwent the same procedure except that no DNA was included in the transfection mixture.

Six hours post-transfection, the medium was replaced with complete DMEM Glutamax-I either supplemented with 0.3 mM NiCl_2_ (#339350, Sigma Aldrich), or only fresh DMEM-Glutamax-I for the non-pathogenic repeat-expressing cells without NiCl_2_ condition. The following day, after confirming transfection by vector GFP fluorescence using ZOE Fluorescent Cell Imager (Bio-Rad), cells were trypsinized (#15050065, Gibco, Thermofisher Scientific) and 5,000 cells were plated on silicon nitride (Si_3_N_4_) membranes (5 mm × 5 mm, 1.5 mm² membrane area, 500 nm membrane thickness, and 200 μm frame thickness, Silson Ltd.) pre-coated with 0.01% poly-L-lysine (#P4707, Sigma Aldrich) and incubated at 37°C, 5% CO_2_ for 48h.

### Cryogenic sample preparation for synchrotron-based X-ray fluorescence microscopy

Approximately 48 hours post-transfection, cells on Si_3_Ni_4_ membranes were briefly washed with 160 mM ammonium acetate solution (#09691-100ML, Sigma Aldrich) freshly prepared with ultra-trace elemental analysis grade water and adjusted to pH 7.4 and 310 mOsm. This step removed the extracellular inorganic salt contaminants from the culture medium. After rinsing, the membranes were quickly blotted and transferred into a temperature and humidity-controlled environmental chamber (37°C, 90% humidity) of the automatic plunge freezer (EM GP2 Automatic Plunge Freezer, Leica) and vitrified by rapidly plunge-freezing in liquid-ethane (−180°C), to preserve the cell’s structural and chemical composition (43).

### Mapping the transfected GFP-positive cells on the Si_3_N_4_ membranes

Vitrified cell monolayers on membranes were imaged with a cryogenic-fluorescence microscope (Leica EM Thunder) equipped with an HC PL APO ×50/0.9 NA cryo-objective at 88K. The brightfield view and bandpass filter cubes of GFP (*λ* = 525/50 nm) were used and DAPI (emission, *λ* = 460/50 nm) allow to detect cell nuclei labelled with Hoechst 33342 (#R37605, Thermofisher Scientific). The resulting fluorescence map was used to locate GFP-positive (transfected) cells for subsequent synchrotron X-ray fluorescence microscopy (SXRFM) analysis. A complete mosaic of the 1.5 × 1.5 mm Si_3_N_4_ active area, containing vitrified cellular regions of interest, was registered as a Z-stack projection over ∼20 µm for each channel with a Nyquist sampling step of 350 nm. Small Volume Computational Clearance (SVCC) from Leica LAS X THUNDER package was applied for deconvolution and blur reduction. Next, maximum intensity projection was applied using LAS X software (Leica Microsystems).

### SXRFM analysis at the ID16A nano-imaging beamline

Experiments were performed at the European Synchrotron Radiation Facility (ESRF) on nanoprobe beamline ID16A (44). The end station operates under high vacuum and features a cryostage that maintains frozen-hydrated samples at 120 K. Combined use of the LEICA EM-VCM and EM-VCT systems enables effective cryotransfer. In this study, the X-ray beam size was approximately 35 nm (horizontal) × 38 nm (vertical) at an excitation energy of 17.1 keV, with an X-ray flux of 2×10¹¹ photons·s⁻¹. A dwell time of 50 ms pixel^−1^ and a stepsize of 50 nm were chosen to ensure high spatial resolution with a reasonable time of acquisition, and while minimizing radiation damage. Fluorescence from each sample pixel is recorded by two custom multielement Silicon Drift Detectors (SDDs), positioned on opposing sides of the sample at 90° relative to the incident X-ray beam. A multielement SDD (Hitachi Ltd.) and an ARDESIA-16 spectrometer based on a monolithic SDD array were employed. The resulting XRF spectra were fitted on a pixel-by-pixel basis using a dedicated Python script based on the use of the PyMca software (45) package, including corrections for detector dead time and normalization of incident beam variations. Elemental areal mass concentrations were calculated using the Fundamental Parameters approach implemented in PyMca, calibrated against a reference standard (RF7-200-S2371 from AXO, Dresden, Germany) with uniform mass depositions in the ng/mm² range (1–3 atomic layers).

### SXRFM image analysis and element quantifications

SXRFM images were analyzed with ImageJ (46). To quantify nickel (Ni) and iron (Fe) levels, regions of interest (ROIs) were defined to isolate cytoplasmic and nuclear compartments: the cytoplasm was segmented based on the uniformly distributed potassium (K) signal, and the nucleus was defined by the zinc (Zn) signal. Fitted and normalized images yielded pixel intensities corresponding to the concentration of each element (ng/mm²). Mean Fe and Ni intensities within the cytoplasmic and/or nuclear ROIs were compared across experimental groups by one-way ANOVA followed by Tukey’s post hoc test (p < 0.05 considered significant). For colocalization analysis, the ImageJ plug-in ScatterJ (version 1.04) (47) was used to compute Pearson’s correlation coefficients between Ni and Fe within each cell. The resulting coefficients for each experimental condition were compared using a Mann-Whitney test, with p < 0.05 considered statistically significant.

### Predictions for iron binding to NOVA2 and other RBPs

We used the AlphaFold v3.0 server (48) to predict the three-dimensional structure of the NOVA2 protein (UniProt accession code Q9UNW9), focusing on its potential to interact with iron. The structural prediction of protein structure and the potential iron-binding sites were analyzed using the molecular visualization software PyMol (49). The predictive outputs were systematically cross-validated using alternative computational tools specialized in metal-binding site prediction, including MIB2 (50). This integrative in silico strategy facilitated the inference of putative iron-binding interactions involving NOVA2, thereby providing computational support for the experimental design.

### Statistics

Statistical analyses are detailed within the respective sections, including information on the type of test used and the criteria for significance.

## RESULTS

### The AUUUC repeat RNA drives the formation of abnormal aggregates in neural stem cells

A frequent feature of RNAs with nucleotide repeat expansions is the adoption of secondary structures, which lead to the formation of aggregates in cells from affected tissues (8,51). The ATTTC repeat has previously been shown to induce abnormal RNA aggregate formation in human embryonic kidney cells (HEK293T) (7). To determine whether the expression of the pathogenic ATTTC repeat in *DAB1* triggers the formation of abnormal aggregates in neural cells, we overexpressed our previously published vector containing either N(ATTTT)_7_ and N(ATTTT)_139_ sequences or the pathogenic repeat insertion in the configuration of (ATTTT)_57_(ATTTC)_58_(ATTTT)_71_, named Ins(ATTTC)_58_ (7) (**Figure 1A**). Using a O-Me-(GAAAU)_5_ probe predicted to hybridize to the AUUUC repeat RNA, we observed by FISH the formation of abnormal nuclear RNA aggregates in human neural stem cells 48 hours post-transfection of a plasmid expressing the pathogenic Ins(ATTTC)_58_, while no aggregates were detected in cells transfected with the non-pathogenic ATTTT repeat sequences, N(ATTTT)_7_ or N(ATTTT)_139_ (**Figure 1B**).

**Figure 1.**
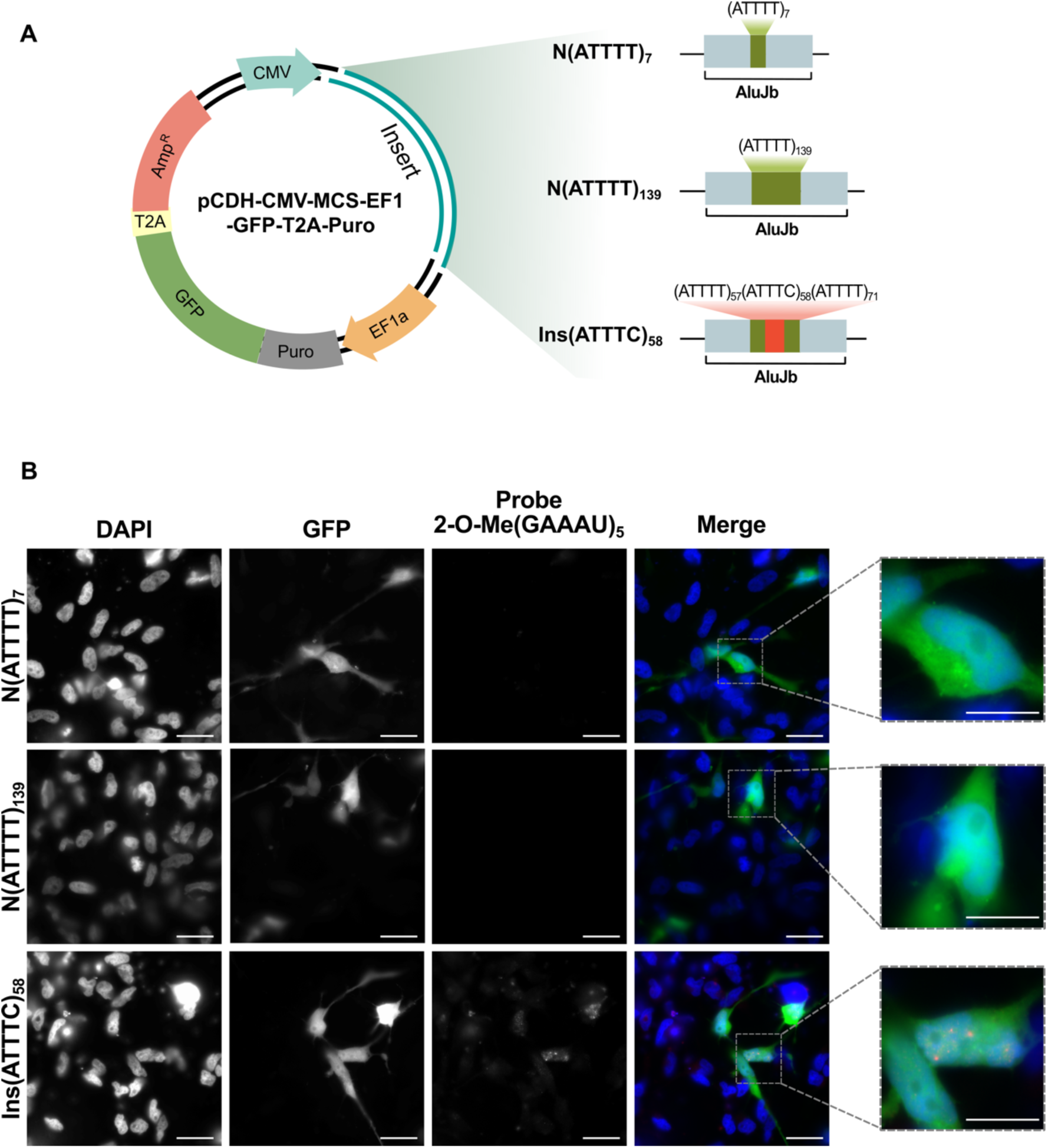
Formation of nuclear AUUUC repeat aggregates in human neural stem cells. **(A)** Schematic representation of the pCDH-CMV-MCS-EF1-GFP-T2A-Puro vector previously described in Seixas et al. (2017), containing three different inserts: a non-pathogenic control harboring (ATTTT)₇ repeats, a longer non-pathogenic (ATTTT)₁₃₉ tract, and a pathogenic repeat configuration consisting of an insertion of (ATTTC)₅₈ repeats embedded within ATTTT tracts, in the configuration (ATTTT)₅₇(ATTTC)₅₈(ATTTT)₇₁, all including the AluJb monomers. **(B)** Overexpression of the pathogenic ATTTC repeat, Ins(ATTTC)_58_, but not the non-pathogenic ATTTT repeats, N(ATTTT)_7_ and N(ATTTT)_139_ in human neural stem cells triggers widespread formation of the aberrant nuclear RNA aggregates, after FISH staining with a probe, 2-O-Me(GAAAU)_5_, predicted to hybridize with the AUUUC repeat, visible 48 hours post-transfection. GFP expression was used as a transfection marker. Images were acquired in Leica DMI6000 FFW inverted epifluorescence microscope, using a HCX PL APO CS 63x/1.30 GLYC 21°C objective. Scale bar = 20 µm.

Thus, these findings demonstrate that the pathogenic AUUUC repeat RNA can drive the formation of abnormal nuclear RNA aggregates in hNSCs.

### Neurodevelopmental RBPs bind to the AUUUC repeat RNA

The sequestration of RBPs by the pathogenic RNA repeat has been described as one of the main mechanisms triggering pathogenicity in non-coding nucleotide repeat expansion diseases (8,52). To identify RBPs binding preferentially to the AUUUC repeat RNA, we performed an RNA pull-down assay using human neuroblastoma cell whole extracts, followed by protein analysis by liquid chromatography-mass spectrometry (LC-MS) (**Figure 2A**). In our screen, the number of RBPs bound to the AUUUC repeat RNA is visibly larger than that bound to the AUUUU repeat RNA (**Figure 2B**). By LC-MS, we identified a total of 28 proteins with higher binding affinity to the pathogenic AUUUC repeat sequence compared with the non-pathogenic AUUUU repeat tract (ANOVA, p<0.001) (**Supp. Table 1**). Those RBPs were classified according to their gene ontology (GO) into categories of biological process, molecular function, cellular components and reactome pathways (**Figure 2C**), revealing enrichment in processes such as RNA processing, mRNA splicing, and mitochondrial RNA catabolism. To prioritize candidates with potential relevance to disease mechanisms, we reviewed the 28 AUUUC-binding RBPs and selected 12 based on two criteria: 1) established roles in neural function (e.g. RNA transport, axonal roles and synaptic function), and/or 2) documented associations with neuromuscular or neurodegenerative diseases in the literature. These 12 proteins are shown in **Figure 3A**. NOVA1 and NOVA2 are alternative splicing factors of neuronal genes known to be implicated in brain disease (53–57). PTBP1/3 are regulators of neuronal splicing crucial for neuronal maturation (58–60). We also detected multiple hnRNPs, including hnRNPA, previously associated with RNA metabolism defects and neurodegeneration (61,62), shown to be sequestered into RNA aggregates in ALS (63), as well as hnRNPR, which is linked to DNA damage repair (64). Matrin-3, implicated in *C9orf72*-ALS (65–67) and nucleolin, known to bind expanded repeats and induce nucleolar stress (31,68) were also pulled down. In addition, we found the ZFR, a regulator of RNA editing at neuromuscular junctions (69), SLIRP, which is essential for mitochondrial function (70) and LRPPRC, a key regulator of mitochondrial activity (71) associated with a neurodegenerative disease (72).

**Figure 2.**
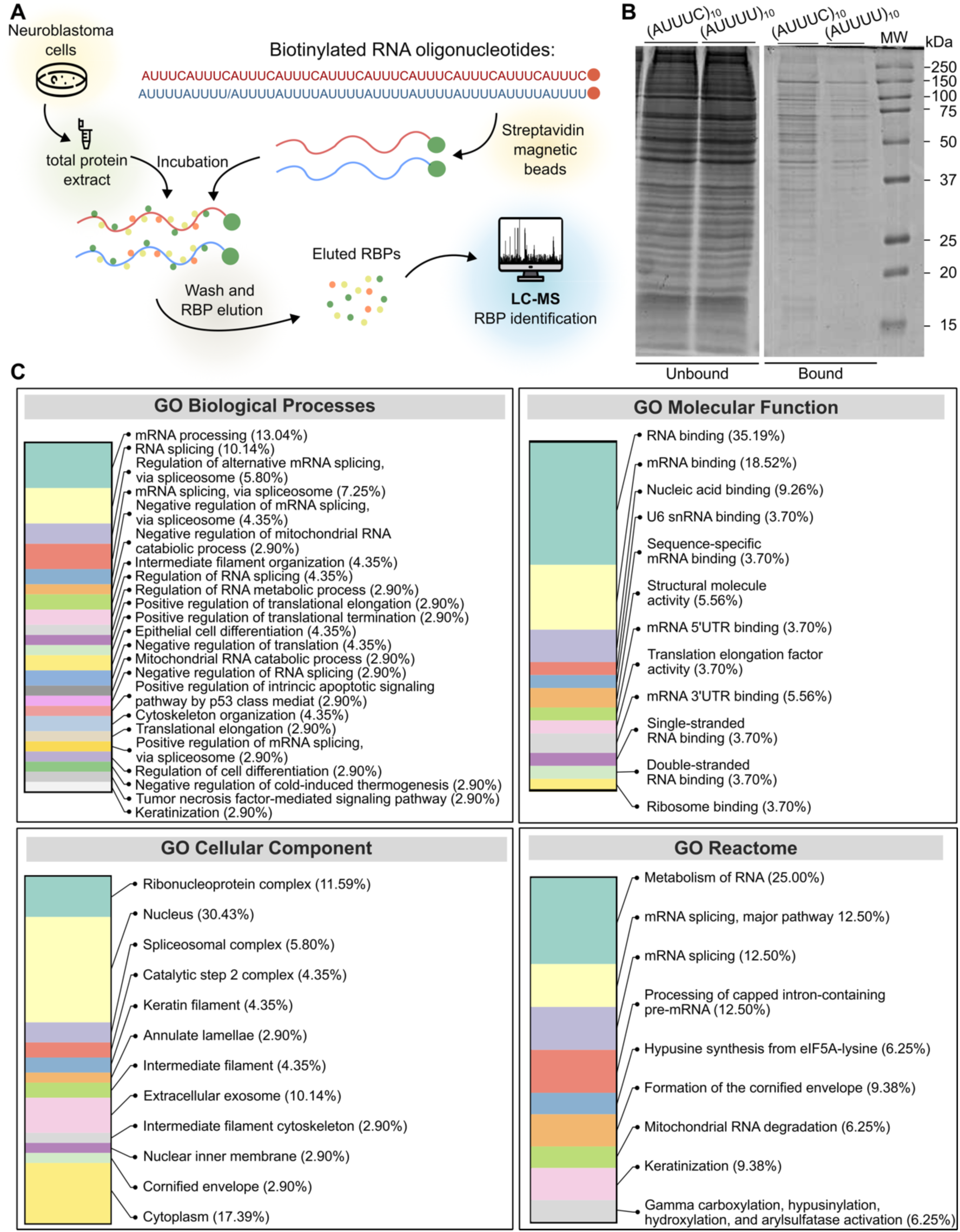
Identification and functional characterization of the RBPs interacting with AUUUC repeat. **(A)** Schematic representation of the experimental workflow in which biotinylated synthesized (AUUUC)₁₀ and (AUUUU)₁₀ RNA molecules were incubated with total protein extracts from neuroblastoma cells. Eluted RBPs were identified via liquid chromatography-mass spectrometry (LC-MS), and the statistical comparisons between the abundance ratio of RBP interaction with the (AUUUC)₁₀ and (AUUUU)₁₀ RNAs were performed using ANOVA. Proteins with a higher abundance ratio in the (AUUUC)₁₀ condition were considered candidate interactors. **(B)** Coomassie-stained SDS-PAGE gel showing proteins bound to biotinylated RNA of either the (AUUUC)_10_ or (AUUUU)_10_ sequences. The (AUUUC)_10_ probe shows visibly more protein bands compared to (AUUUU)_10_, indicating a greater number of RBPs associated with the AUUUC sequence. **(C)** Gene Ontology (GO) enrichment analysis of the 28 proteins that showed increased binding to the (AUUUC)₁₀ repeat compared to the (AUUUU)_10_. GO terms were identified using the Functional Annotation Tool available in the Database for Annotation, Visualization and Integrated Discovery (DAVID), with a significance threshold of adjusted p<0.05.

**Figure 3.**
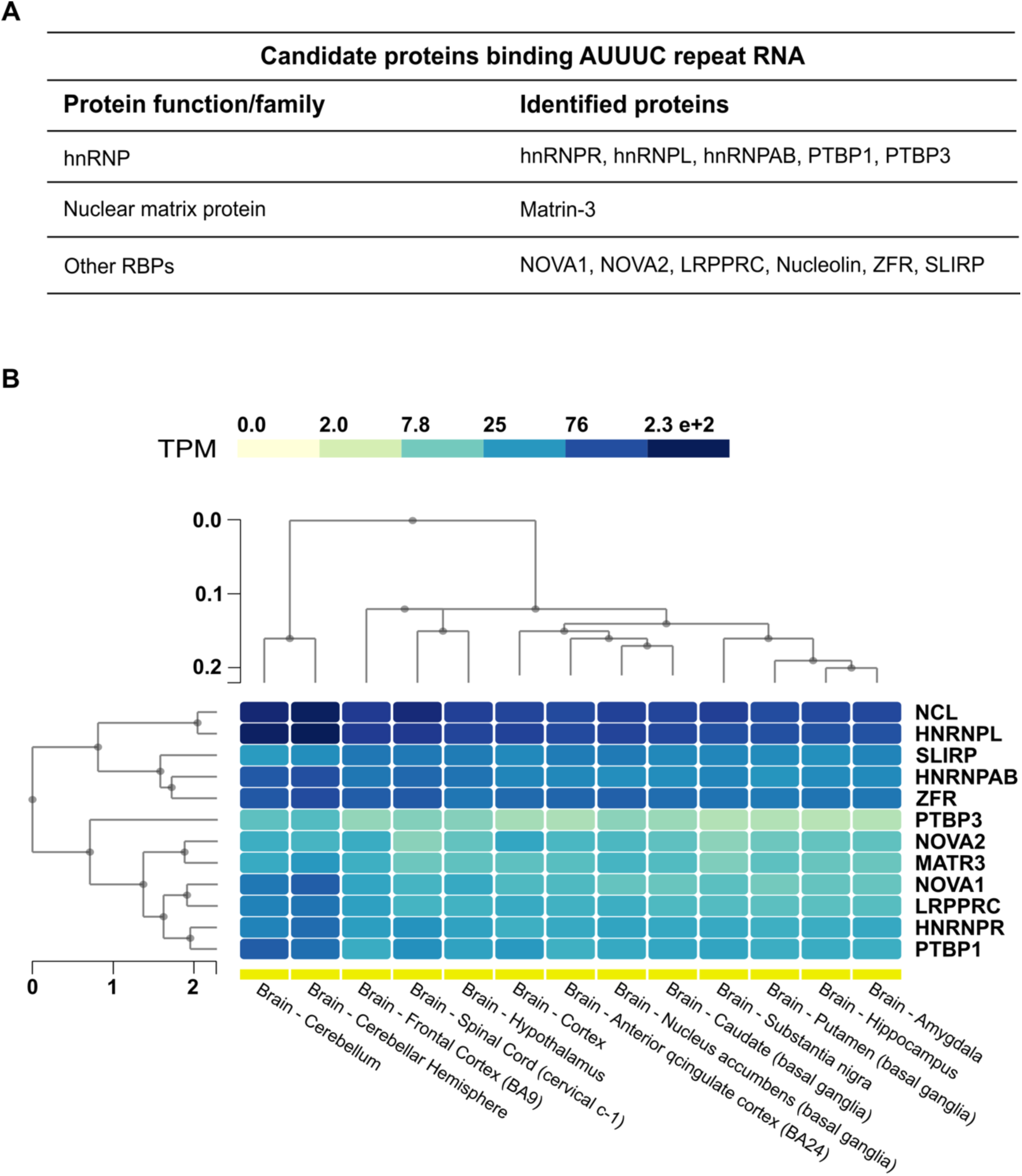
RBPs with neural functions bound to the AUUUC. **(A)** Table listing the top 12 candidate RBPs interacting with (AUUUC)_10_ based on their crucial neural function and/or association with neuromuscular or neurodegenerative diseases, categorized by protein function or family. **(B)** Heatmap showing the expression levels, in transcripts per million (TPM), of the top 12 identified RBPs across various human brain regions, using GTEx RNA-seq data. The color scale represents increasing expression levels from low (light yellow) to high (dark blue), and hierarchical clustering indicates similarities in gene expression profiles across tissues.

To further support their neurological relevance, we constructed a heatmap of their expression profiles across various human brain regions, using GTEx RNA-seq data (**Figure 3B**), showing that these RBPs are broadly and highly expressed in brain tissue, consistent with a potential role in nervous system function and disease.

Our findings provide evidence that the AUUUC repeat sequence binds RBPs essential for brain development and neuronal differentiation, suggesting that disruption of their role may contribute to neurodevelopmental and neurodegenerative diseases.

### NOVA2 colocalizes with the AUUUC repeat RNA aggregates in neural stem cells

NOVA2 is an RBP known for its pivotal role in regulating hundreds of alternative splicing events in neurons during neurodevelopment, fundamental for neuronal migration and brain development (53–55,73). The predicted RNA-binding motif for NOVA2 is YCAY, where Y represents a pyrimidine (74). This motif is present in the AUUUC repeat tract, only intercalated by a pyrimidine nucleotide in the pathogenic repeat insertion. Moreover, our RNA pull-down experiment revealed a higher affinity of NOVA2 for the pathogenic AUUUC repeat RNA compared with the non-pathogenic AUUUU repeat tract, and interestingly, we observed motor neuron defects caused by the injection of the pathogenic AUUUC repeat RNA during embryonic development in our transient zebrafish model (75). Thus, we hypothesized that NOVA2 is sequestered by AUUUC repeat RNA to form abnormal aggregates in hNSC. Since both *DAB1* and *NOVA2* are highly co-expressed in brain tissues (**Supp. Figure 1**), the colocalization of NOVA2 with AUUUC repeat RNA aggregates observed in neural stem cells is biologically conceivable. To test this, we co-transfected hNSC with our published mammalian expression vectors (7) expressing either the pathogenic or the non-pathogenic RNAs embedded in their flanking sequences (**Figure 1A**), and a plasmid encoding NOVA2 fused at its C-terminus to a histidine tag, NOVA2-His-tag, for detection by an anti-His antibody (**Figure 4A**). Using a combination of RNA-FISH and immunofluorescence, we observed by confocal microscopy that NOVA2 colocalizes with the abnormal AUUUC repeat RNA aggregates in multiple cells expressing the pathogenic repeat but not in non-pathogenic repeat control cells (**Figure 4B**). We did not observe the colocalization of AUUUC repeat RNA and NOVA2 in all aggregates, showing that the sequestration is sometimes a dynamic process, instead of irreversible sequestration with the total compromise of protein function, as usually observed for the interaction of other proteins with pathogenic repeat RNAs like in *C9orf72*-ALS (76). The AUUUC repeat RNA aggregates were predominantly nuclear, as previously demonstrated in non-neural cells (7). In hNSC, NOVA2 was mostly located in the perinuclear region in cells expressing the non-pathogenic repeat, while a shift for the nuclear core was observed in cells expressing the pathogenic repeat. We, thus, hypothesized that NOVA2 could be mislocalized in hNSCs expressing the SCA37 repeat, due to its sequestration in the toxic AUUUC repeat RNA aggregates, when compared with cells transfected with the non-pathogenic repeat. To explore this, we quantified the intensity of NOVA2 immunofluorescence in the periphery and core regions of nuclei from cells belonging to the different conditions (**Supp. Figure 2**). Although the results obtained suggest a tendency for NOVA2 to exhibit a higher signal at the periphery of nuclei in cells with the shorter non-pathogenic repeat, compared with those expressing the pathogenic repeat, the differences were not statistically significant.

**Figure 4.**
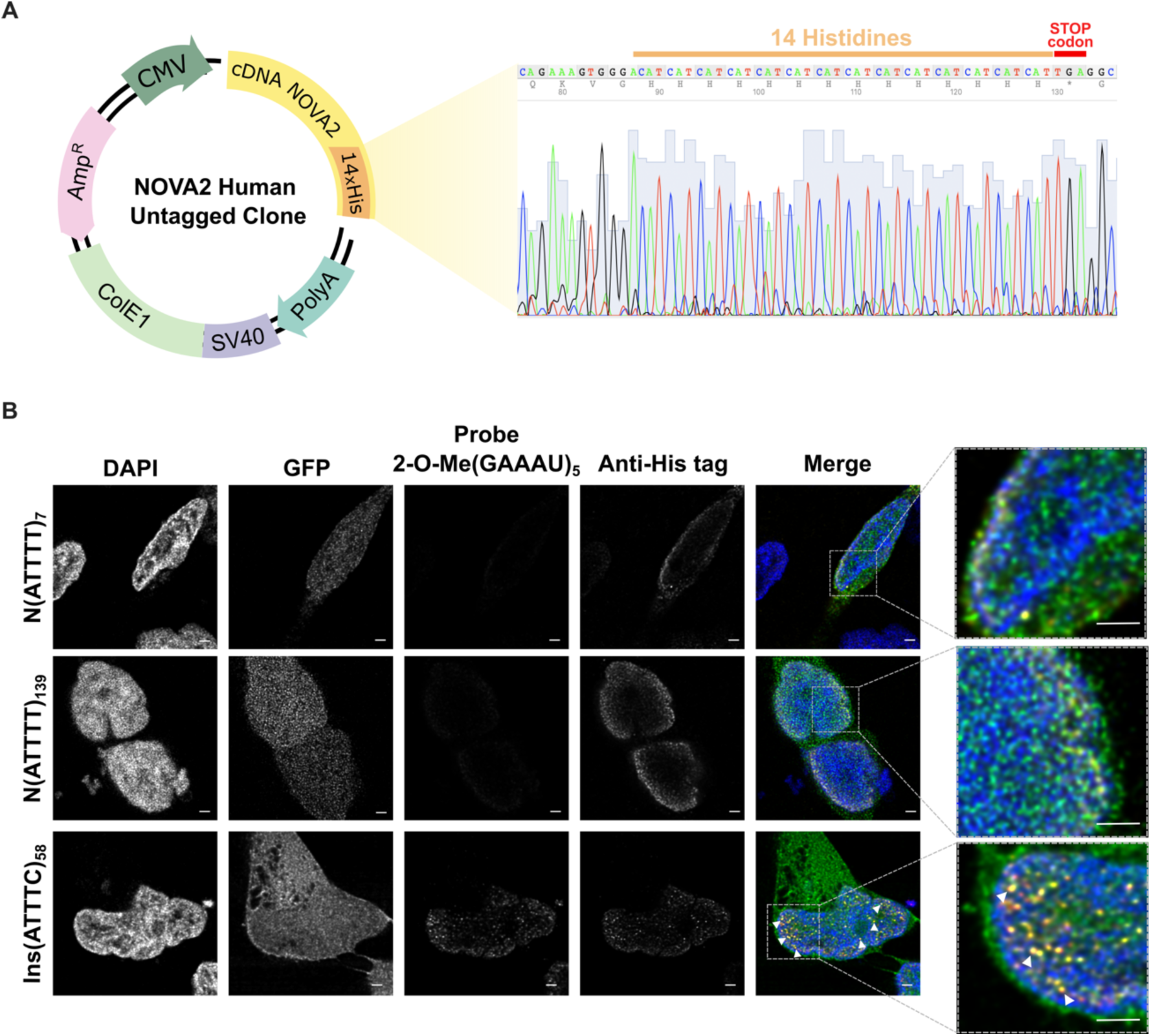
NOVA2 colocalizes with nuclear AUUUC repeat aggregates in human neural stem cells. **(A)** Schematic representation of the NOVA2 Human Untagged Clone (#SC303210, OriGene) containing the full-length NOVA2 cDNA (NM_002516) that we modified by inserting a 14×His-tag in-frame at the C-terminus of the NOVA2 open reading frame. The scheme illustrates the NOVA2 coding sequence (yellow), the inserted 14×His-tag (orange), followed by the stop codon (red). **(B)** hNSC co-overexpressing the non-pathogenic N(ATTTT)_7_, N(ATTTT)_139_ or the pathogenic Ins(ATTTC)_58_ sequences and NOVA2-His-tag, show aberrant nuclear AUUUC repeat aggregates that colocalize with NOVA2 (white arrows), but not AUUUU repeat accumulation, after staining with a specific FISH probe followed by immunofluorescence with the Anti-his-tag antibody, visible at 48 hours post-transfection. GFP expression was used as a marker for transfection. Images acquired in Leica Microsystems TCS SP5 II confocal microscope, using a HC PL APO Lbl. Blue 63× /1.40 Oil objective with 8× zoom. Scale bar = 2 µm.

Our results provide the first evidence that the NOVA2 protein is sequestered within the abnormal AUUUC repeat RNA aggregates and with a tendency for mislocalization to the nuclear core.

### Iron accumulates in the AUUUC repeat RNA-NOVA2 aggregates

Iron homeostasis is critical for the survival of neural stem cells and influences pathology during neuronal ageing (77). Motivated by the emerging connection between iron dysregulation and neurodegenerative disorders (78–80), we investigated iron dyshomeostasis in SCA37, using high-resolution SXRFM imaging analysis. We used this technique to map and localize NOVA2 intracellularly, based on the high affinity of nickel for the His-tag cloned at the C-terminal of the protein, and simultaneously analyze the distribution and concentration of a variety of trace elements. Before SXRFM analysis, we seeded HEK293T cells co-transfected with the vectors expressing either the non-pathogenic ATTTT repeat or the pathogenic ATTTC repeat and the NOVA2-His-tag plasmid onto Si_3_N_4_ membranes. We vitrified the cells forty-eight hours post-transfection to ensure the preservation of cellular components, and we mapped the distribution of GFP-positive cells adhered to the membrane by analyzing the frozen-hydrated cells in their near-native state, by wide-field fluorescence cryo-microscopy (**Supp. Figure 3A-B**). This guaranteed that subsequent observations under cryogenic conditions with the synchrotron X-ray fluorescence microscope were focused only on RNA-expressing cells.

To evaluate the cellular vitrification quality at specific positions, we acquired elemental distribution XRF maps of the entire cells. We performed initial low-resolution coarse scans (400 nm × 400 nm, 100 ms exposure), a few minutes long. In cells with a homogeneous distribution of potassium, which confirmed integrity, we acquired high-resolution nanoscopic images (continuous fast scan at 50 nm stepsize) for the different cellular experimental conditions. Subsequently, we examined in greater detail our regions of interest, specifically iron- and nickel-rich hotspots, by acquiring ultra-high-resolution scans (20 nm stepsize) to analyze fine structural and compositional details. Across all conditions, we acquired 24 coarse scans (400 nm resolution), 24 fine scans (50 nm resolution) (**Supp. Figure 4**) and 14 ultra-high-resolution maps (20 nm resolution), focusing on iron-rich nanoscopic aggregates of RNA-expressing GFP-positive cells (**Supp. Figure 5**). We found, as expected, a higher concentration of zinc (Zn) within the nucleus, in agreement with previous findings (81), consistent with its role in nuclear processes. Thus, Zn distribution maps were used to identify and delineate the nuclear region. Potassium (K), known to be uniformly distributed across the cell, served as a marker for overall cell morphology and contours. By combining these elemental maps, the nucleus and cytoplasm were distinguished, enhancing the interpretation of cellular structure and composition **(Figure 5)**.

**Figure 5.**
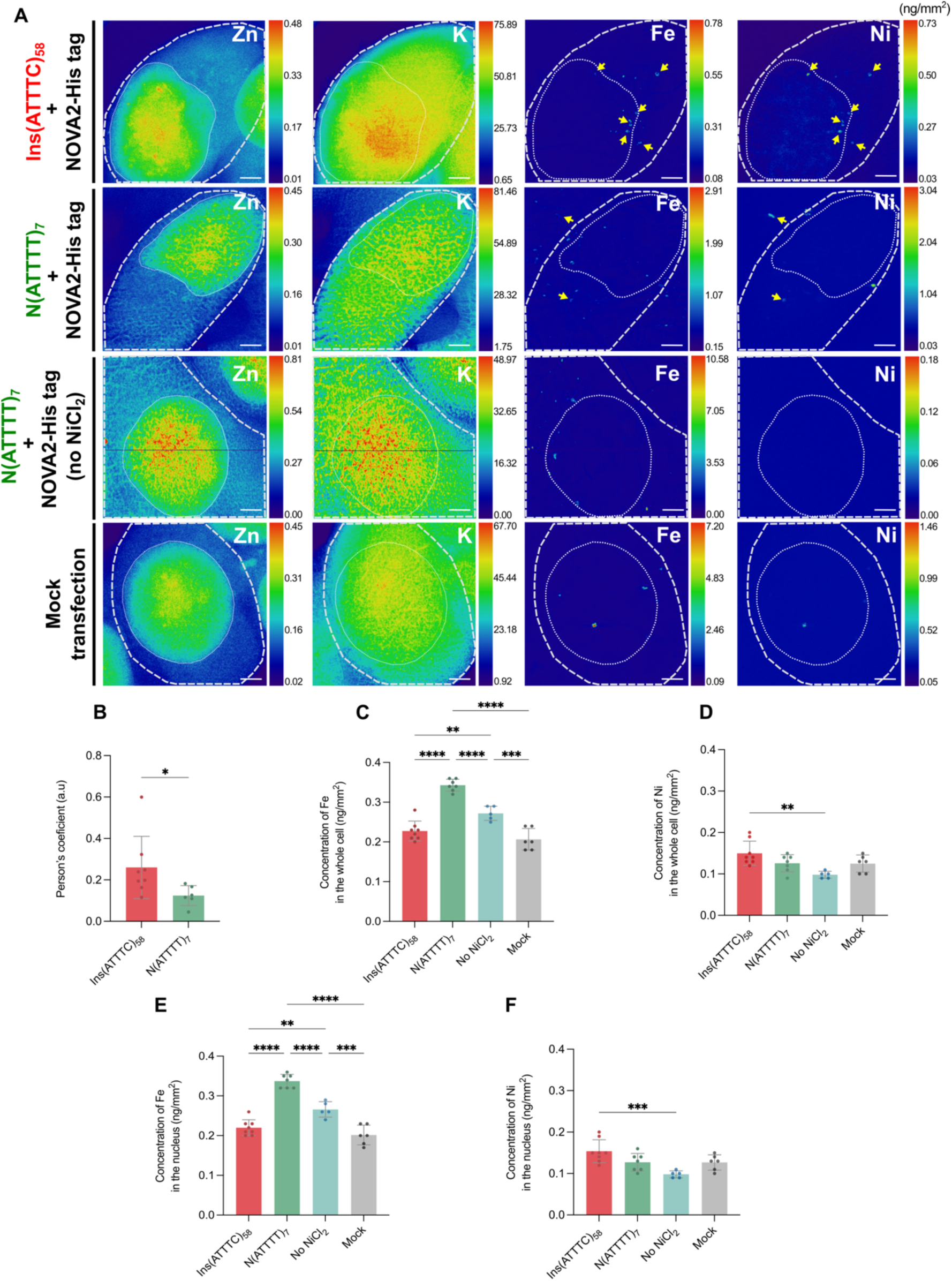
Iron colocalizes with NOVA2 in the synchrotron X-Ray fluorescence analysis. **(A)** Representative high-quality scans of HEK293T cells were analyzed under each experimental condition: (1) cells co-expressing either the SCA37 pathogenic repeat insertion, Ins(ATTTC)_58_, or the non-pathogenic N(ATTTT)_7_ and NOVA2-His-tag; (2) cells expressing the non-pathogenic N(ATTTT)_7_ and NOVA2-His-tag sequence, but with no NiCl_2_ supplementation; and (3) mock-transfected cells (expressing no plasmids). The fluorescence images highlight the localization and expression patterns of iron (Fe) and nickel (Ni) under each condition. Zinc (Zn) delineates the nucleus and potassium (K) the cytoplasm. The X-Ray beam size was approximately 35 nm (horizontal) × 38 nm (vertical) at an excitation energy of 17.1 keV, with an X-Ray flux of 2×10^11^ photons.s^-1^. A dwell time of 50 ms.pixel^−1^ and a stepsize of 50 nm were used. **(B)** Pearson’s Correlation Coefficient for iron and nickel distribution was calculated for individual cells under each experimental condition. Statistical significance between the conditions was determined using a Mann-Whitney test, with a threshold of p<0.05 for significance. **(C-F)** Graphical representation of the concentration of **(C)** iron and **(D)** nickel, in the whole cell; and concentration of **(E)** iron and **(F)** nickel, in the nucleus. For (C), (D), (E) and (F), statistical significance was determined using One-Way ANOVA, with p<0.05 considered significant. Scale bar = 50 pixels.

We found a significantly higher co-occurrence of iron and nickel signals in cells expressing the SCA37 pathogenic repeat compared with cells with the non-pathogenic repeat **(Figure 5)**. To assess colocalization, we constructed scatterplots comparing iron and nickel signal overlay in cells from both conditions (**Supp. Figure 6**) and calculated the Pearson’s Correlation Coefficient for iron and nickel distribution for individual cells (**Figure 5B**). We observed a significantly higher Pearson’s Correlation Coefficient in cells expressing the pathogenic repeat compared with cells presenting the non-pathogenic repeat, indicating increased colocalization (Mann-Whitney test, p < 0.05). Most of the Fe/Ni colocalization signals observed in cells with the pathogenic AUUUC repeat RNA were within the nuclear region. However, cytoplasmic signals were also detected, and they may result from the rupture of the nuclear membrane during cell division, given the high proliferation rate of embryonic kidney cells. We detected a few colocalized lower signals of iron and nickel in the cells with the non-pathogenic repeat, which can be explained by the overexpression of both AUUUU repeat RNA and NOVA2-His-tagged protein, which may also lead to the production of smaller protein aggregates capable of accumulating iron, though at a much lower frequency and quantity. Interestingly, when analyzing the iron-rich structures in detail (**Supp. Figure 5**), we found no overlap between iron and sulfur elemental maps, meaning that those are not Fe/S clusters, structures with an important role in electron transfer during mitochondrial respiration commonly found in cells (82). Additionally, observations of the iron-rich nanoscopic aggregates revealed distinct differences in morphology between the conditions. In cells transfected with the SCA37 pathogenic repeat, the Fe/Ni structures appear more diffuse (**Supp. Figure 5A**), whereas in cells transfected with the non-pathogenic repeat, these structures exhibit a more rounded shape (**Supp. Figure 5B)**. This rounded morphology may suggest the presence of metallic particles, while the diffuse structures found in the SCA37-transfected cells might result from the presence of cellular structures, like RNA/protein aggregates, which supports our hypothesis. Moreover, it is important to note that although the rapid rinsing with ammonium acetate buffer may not remove all contaminants, the vitrification of cells by plunge freezing preserves their chemical integrity and maintains them in a near-native state. Therefore, it is highly unlikely that the observed intracellular colocalization of iron and nickel in cells transfected with the pathogenic repeat is an artefact of sample preparation.

These results show that the AUUUC repeat RNA-NOVA2 aggregates accumulate iron, suggesting an iron imbalance triggered by the SCA37 pathogenic repeat.

### Lower free iron content in SCA37-expressing cells and iron dyshomeostasis

Given the increasing recognition of iron dyshomeostasis as a contributor to neurodegeneration, we explored whether the pathogenic SCA37 repeat might affect intracellular iron balance. We quantified intracellular iron and nickel levels in cells, measuring their intensities across the whole cell and nucleus (**Figure 5C-F**). We found a significantly lower iron concentration in cells expressing the SCA37 repeat when compared with cells presenting the non-pathogenic repeat (**Figure 5C** and **5E**; One-Way ANOVA, p<0.05). Regarding nickel concentration, because of the sensitivity of the method, measurements from cells transfected with the non-pathogenic ATTTT repeat cultured without NiCl_2_ supplementation in the medium provided a baseline (background) nickel level. Since nickel is a trace element with physiologically low concentrations in mammalian cells, only small differences were expected across conditions supplemented with 0.3 mM NiCl_2_. Nevertheless, our results showed that the cells expressing the SCA37 repeat sequence exhibited significantly higher nickel concentrations compared with that carrying the non-pathogenic repeat tract in the absence of NiCl_2_ supplementation (**Figure 5D** and **5F**; One-Way ANOVA, p<0.05), while no significant differences were observed among the ATTTT repeat-transfected cells, mock-transfected cells and those without NiCl_2_ supplementation (**Figure 5D** and **5F**; One-Way ANOVA, p>0.05). The observed higher nickel concentrations in cells transfected with the SCA37 repeat strongly suggest that this pathogenic repeat forms aggregates capable of binding NOVA2, leading to its retention.

In parallel, to investigate whether NOVA2 could directly interact with iron, we used the AlphaFold3 server (48) to predict metal-binding residues in NOVA2 (**Supp. Figure 7A**). Despite moderate overall confidence for the model, the predicted iron-binding region in NOVA2 exhibited higher confidence scores, suggesting a plausible interaction site. Additionally, the predicted geometry and distances (∼2.8 Å) matched typical iron-binding motifs and were corroborated by MIB2 predictions (**Supp. Figure 7B**), supporting NOVA2 potential role in iron coordination.

Together, these results showed that the SCA37 repeat has the capacity to disrupt iron homeostasis, giving new insights into the cellular consequences of DNA/RNA repeat expansions.

## DISCUSSION

In this study, we show that the pathogenic AUUUC repeat RNA sequesters NOVA2, a neuronal splicing regulator, into the abnormal RNA aggregates in neural cells. We also demonstrate that these RNA aggregates accumulate iron, leading to cellular alterations in iron content. As NOVA2 and iron colocalize in cells, we propose that SCA37 pathogenesis may involve an AUUUC repeat RNA gain-of-function by imprisoning NOVA2 and likely other RBPs, leading to the perturbation of iron homeostasis, a process essential for neuronal viability.

This work shows the critical role of DNA/RNA repeats in neural cells. NOVA2 is a neural-specific RBP that regulates hundreds of alternative splicing events in the brain (57) and plays an essential role in neuronal development, axon pathfinding and guidance, as evidenced by the *Nova2* knockout mice, which exhibit severe defects in corpus callosum formation and motor neuron axon outgrowth (53). A partial loss of NOVA2 function impairs its ability to regulate neurite outgrowth and axonal guidance (55). Moreover, NOVA2 has also been shown to regulate neuronal migration through the alternative splicing of the *Dab1* gene, a key mediator of reelin signalling (73). Although the sequestration of RBPs in RNA foci does not fully indicate complete functional inactivation or irreversible sequestration (76), it is likely these proteins have their function, at least, partially compromised. Supporting NOVA2 loss-of-function, our CAGE data from SCA37 patient-derived induced pluripotent stem cells (iPSCs) reveals dysregulation of NOVA2 splicing targets, including *DAB1* (83). Remarkably, in our transient zebrafish model of SCA37, motor neuron defects caused by the injection of the pathogenic AUUUC repeat RNA during embryonic development can be rescued through co-injection of NOVA2 (75). Together, these findings suggest that AUUUC-mediated toxicity may involve the sequestration of NOVA2 and additional AUUUC-binding RBPs into the RNA aggregates. However, the downstream effect of this sequestration, its subsequent loss-of-function and how it may play a critical role in SCA37 neuropathology remain to be determined.

These results also demonstrate a crucial role of DNA/RNA repeats in iron homeostasis. Iron homeostasis has been shown to modulate alternative splicing of genes encoding proteins promoting or inhibiting apoptosis (84). Iron has a direct effect on RNA recognition by the splicing factor SRSF7, in which the level of iron influences the binding of this factor to the alternative splicing target region, regulating apoptosis (84). While the modulation of the alternative splicing function of NOVA2 by iron has not been described, our results raise the hypothesis of a potential role for iron in NOVA2 activity. Notably, MRI studies in SCA3, an identical disease with abnormal protein aggregation in neurons, have revealed increased brain iron deposition as an early event in disease pathogenesis (85). In this context, our observation of iron accumulation within the abnormal AUUUC repeat RNA aggregates further supports the involvement of iron dyshomeostasis in SCA37, similarly contributing to neurodegeneration.

X-ray fluorescence (XRF) imaging has previously been used to investigate iron distribution and metal concentrations in Friedreich’s ataxia (FRDA), characterized by the bi-allelic expansion of a GAA repeat in the intronic region of the FXN encoding frataxin, leading to lower iron-chaperone frataxin levels (86). Nanoscopic XRF analysis of cryofrozen FRDA fibroblasts revealed iron-rich clusters in the cytoplasm and a reduction in zinc levels (87). Further, the authors analyzed freeze-dried FRDA fibroblasts and identified iron-rich organelles, including the endoplasmic reticulum, mitochondria, and lysosomes, as well as nanoscopic iron hot spots within the cytosol (78). Iron dysregulation is a hallmark of the repeat expansion disease Friedreich’s ataxia (79) and other neurodegenerative disorders such as Alzheimer’s, Parkinson’s, and ALS (80). In FRDA, frataxin deficiency disrupts iron-sulfur cluster biogenesis and causes mitochondrial iron accumulation (79). Similarly, in Alzheimer’s, Parkinson’s, and ALS, dysregulated iron metabolism contributes to neurotoxicity and oxidative stress (80). Moreover, it is known that aggregates of the RBP TDP-43, a hallmark of ALS, are also observed in disorders grouped as neurodegeneration with brain iron accumulation (88), showing a link between RBP aggregation and iron accumulation. The mutant ALS protein, SOD1, promotes modifications that drive TDP-43 mislocalization and aggregation (89), and considering the present study, it is of interest to note that in SOD1-mutant mice, treatment with iron chelators has been found to attenuate TDP-43 aggregation, whereas vehicle-treated animals display cytoplasmic TDP-43 inclusions in motor neurons (90). In spite of that, until now, no other study has shown the accumulation of iron in RNA aggregates containing RBPs.

This study marks the first application of a novel SXRFM approach for the precise localization of proteins in near physiological conditions while simultaneously mapping elemental distributions. Using a His-tagged NOVA2 construct, we have tracked its sequestration and colocalization with iron within the AUUUC repeat RNA aggregates. Given the limited beamtime access and the fact that it is the first application of SXRFM to RNA aggregate biology, we used HEK293T to maximize the chance of obtaining interpretable data. The use of HEK293T cells serves as a proof-of-concept platform for applying SXRFM to repeat-associated RNA biology. While there is still room for improvement, our method shows great promise, and future work may be done to extend these observations to neural models now that the technique has been validated. Indeed, this method uses cells vitrified and maintained in a frozen-hydrated state for analysis under cryogenic conditions, thereby ensuring excellent preservation of the cellular structure and intracellular chemical integrity. Moreover, its high sensitivity at the nanoscale resolution (20-50 nm) and the ability to detect over 15 trace elements simultaneously, provide a powerful platform for mapping protein-metal interactions *in situ*. With further optimization, this technique can be adapted to study multiple tagged proteins concurrently with different metal-affinity markers, broadening its applicability and facilitating the simultaneous study of multiple interacting proteins.

These results strongly suggest that both the loss of RBP function and iron dyshomeostasis may be key contributors to SCA37 pathology. Several unanswered questions and intriguing directions for future research have emerged from this study. In particular, NOVA2 and the remaining 11 neural RBPs identified in the RNA pull-down will need to be functionally investigated to determine whether their target transcripts are dysregulated, mis-spliced, or have altered expression. Moreover, further investigation into iron dyshomeostasis in neurodegenerative repeat expansion disorders is fundamental to understanding disease mechanisms and uncovering novel therapeutic targets.

In conclusion, these findings provide evidence, for the first time, that a pathogenic RNA, the pathogenic SCA37 repeat, sequesters essential RBPs, such as NOVA2, influencing the intracellular dynamics of iron and contributing to iron dyshomeostasis.

## ACKNOWLEGDMENTS

We are grateful to Dr. Pereira-Castro for comments on the manuscript. We acknowledge Dr. Kenya Ishikawa for the advice on the RNA pull-down protocol. We are thankful to Drs. Pedro Pereira and Joana Santos for technical assistance with the IVA cloning protocol. We thank Dr. Samira Acajjaoui for technical assistance with the cell culture preparation for the SXRFM experiment. We acknowledge the European Synchrotron Radiation Facility (ESRF) for the provision of synchrotron radiation facilities at ID16A beamline in the frame of proposal LS-3136 and the CIBB lab for cell culture preparation. Data from ID16A sessions correspond to the DOI https://doi.org/10.15151/ESRF-ES-1716285807.

This work was supported by Fundo Europeu de Desenvolvimento Regional (FEDER), through the COMPETE 2020 Operational Program for Competitiveness and Internationalization (POCI) of Portugal 2020, FCT and Ministério da Ciência, Tecnologia e Ensino Superior (Portugal), in the framework of the project POCI-01-0145-FEDER-029255 (PTDC/MED-GEN/29255/2017) to I.S and J.B; and by COMPETE2030-FEDER-00691000, Grant No. 15801, funded by COMPETE2030 and FCT under MPr-2023-12 to I.S. AS.F is a recipient of the FCT PhD scholarship 2021.05757.BD.

## AUTHOR CONTRIBUTIONS

A.S.F conceived of this project with J.R.L, H.O, J.B, S.B, S.M.R and I.S, led the development and optimization of the cellular experiments, performed the experiments, analyzed and interpreted data, wrote the paper and produced the figures. J.R.L conceived of the project with A.S.F, H.O, J.B, S.B, S.M.R and I.S, performed the RNA-pull down experiment, and reviewed the manuscript. P.S developed the macro for NOVA2 localization analyses, participated in the microscopy image acquisition and reviewed the manuscript. H.O conceived of this project with A.S.F, J.R.L, J.B, S.B, S.M.R and I.S, performed the LC-MS experiments, analyzed the data and reviewed the manuscript. M.S.L supported the cell culture of SXRFM experimental design and procedures and reviewed the manuscript. J.B conceived of this project with A.S.F, J.R.L, H.O, S.B, S.M.R and I.S and reviewed the manuscript. S.B conceived of this project with A.S.F, J.R.L, H.O, J.B, S.M.R and I.S, designed and performed the SXRFM experiments, image acquisition and processing, oversaw the SXRFM analysis and reviewed the manuscript. S.M.R conceived of this project with A.S.F, J.R.L, H.O, J.B, S.B, and I.S, oversaw the SXRFM experiments and analysis, and reviewed the manuscript. I.S conceived of this project with A.S.F, J.R.L, H.O, J.B, S.B, and S.M.R and oversaw all the analysis and experiments performed and reviewed the manuscript.

