## Supplementary Figures and Tables for "The AUUUC repeat RNA aggregates sequester RNA-binding proteins like NOVA2 and lead to iron dyshomeostasis in spinocerebellar ataxia type 37"

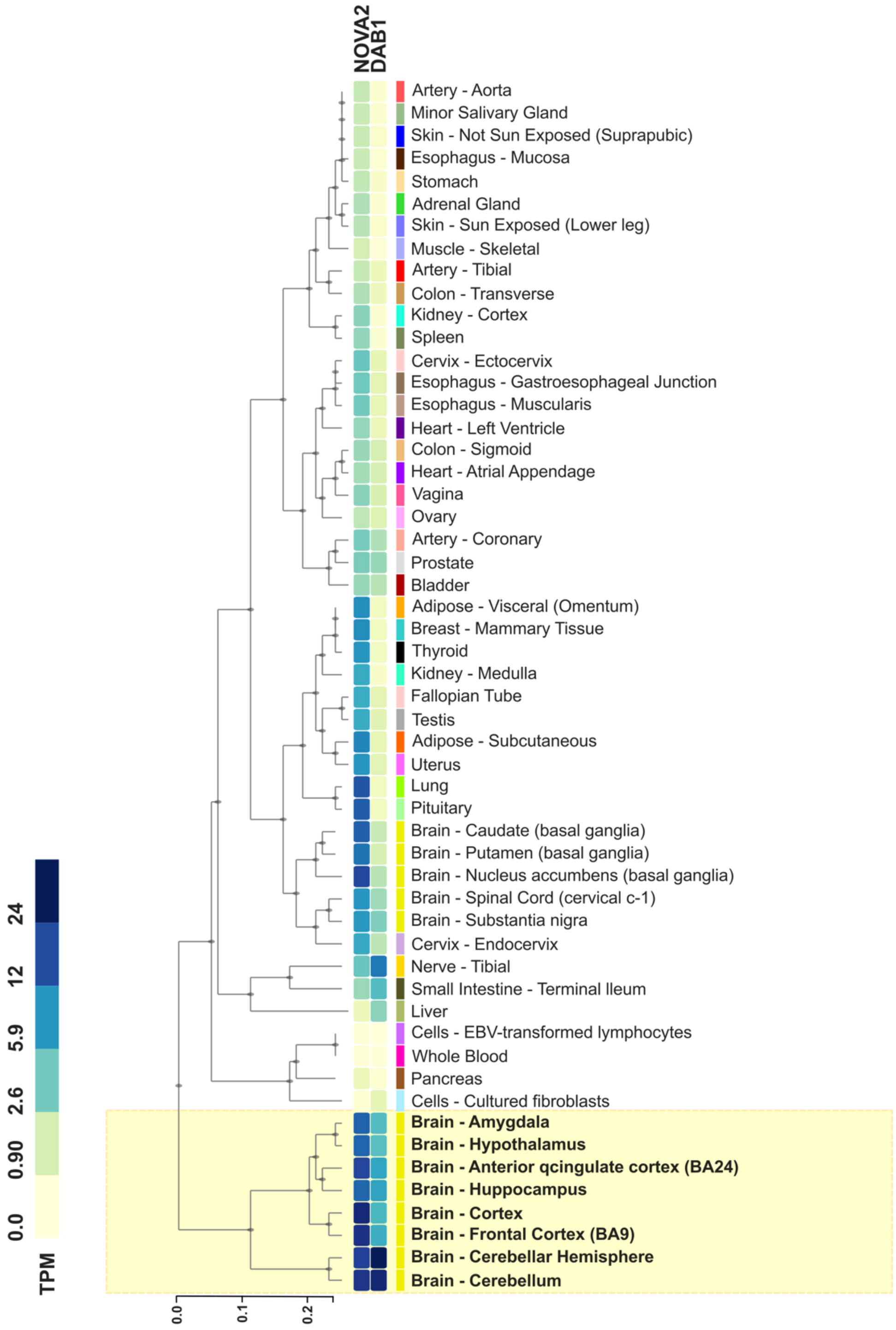


**Supplementary Figure 1 – Co-expression of *DAB1* and *NOVA2* in brain tissues.** The heatmap shows transcript expression levels, in TPM, of *DAB1* and *NOVA2* across various human tissues using data from the GTEx project. Hierarchical clustering reveals a clear pattern of brain-specific co-expression of both genes. Notably, high expression levels are observed in multiple brain regions, including the cerebellum, highlighted in the yellow box. This co-expression supports the sequestration of NOVA2 by the transcribed AUUUC in neural tissues.

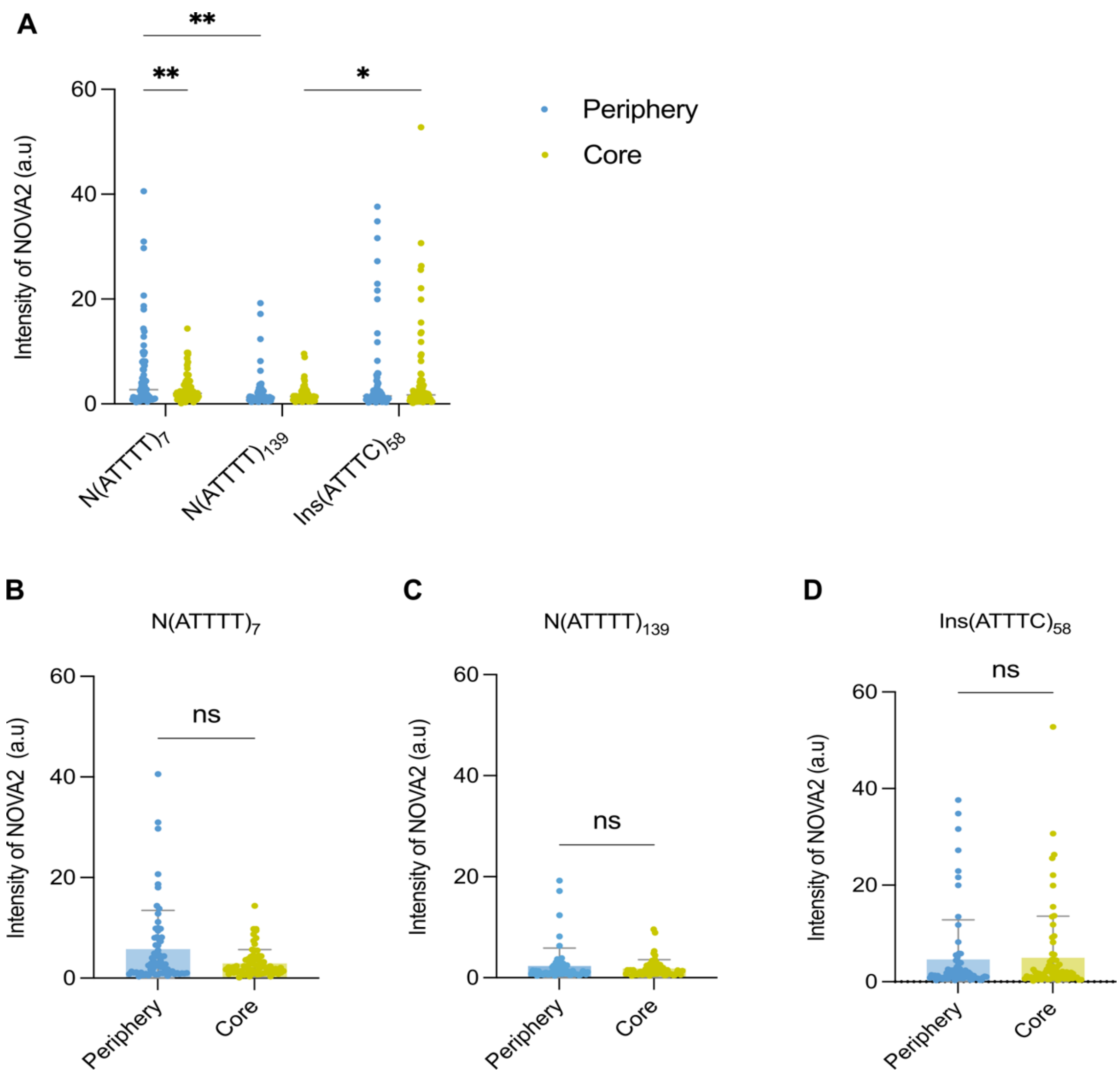


**Supplementary Figure 2 - NOVA2 distribution in human neural stem cells co-expressing pathogenic or non-pathogenic repeats.** hNSC co-expressing the non-pathogenic N(ATTTT)$_7$, N(ATTTT)$_{139}$ or the SCA37 Ins(ATTTC)$_{58}$ repeats and NOVA2-His tag were analyzed in a total of 69 cells for N(ATTTT)$_7$, 61 cells for N(ATTTT)$_{139}$ and 74 cells for Ins(ATTTC)$_{58}$. **(A)** Graphical representation of NOVA2 intensity in the periphery and core of the cells. Statistical significance between the conditions was determined using a 2-way ANOVA test, with a threshold of $p<0.05$ for significance. Graphical representation of NOVA2 intensity, both in the periphery and core of the cells, for **(B)** N(ATTTT)$_7$ transfected cells; **(C)** N(ATTTT)$_{139}$ transfected cells and **(D)** Ins(ATTTC)$_{58}$ transfected cells. Statistical significance between the conditions of (B), (C) and (D) were determined using a Mann-Whitney test, with a threshold of $p<0.05$ for significance.

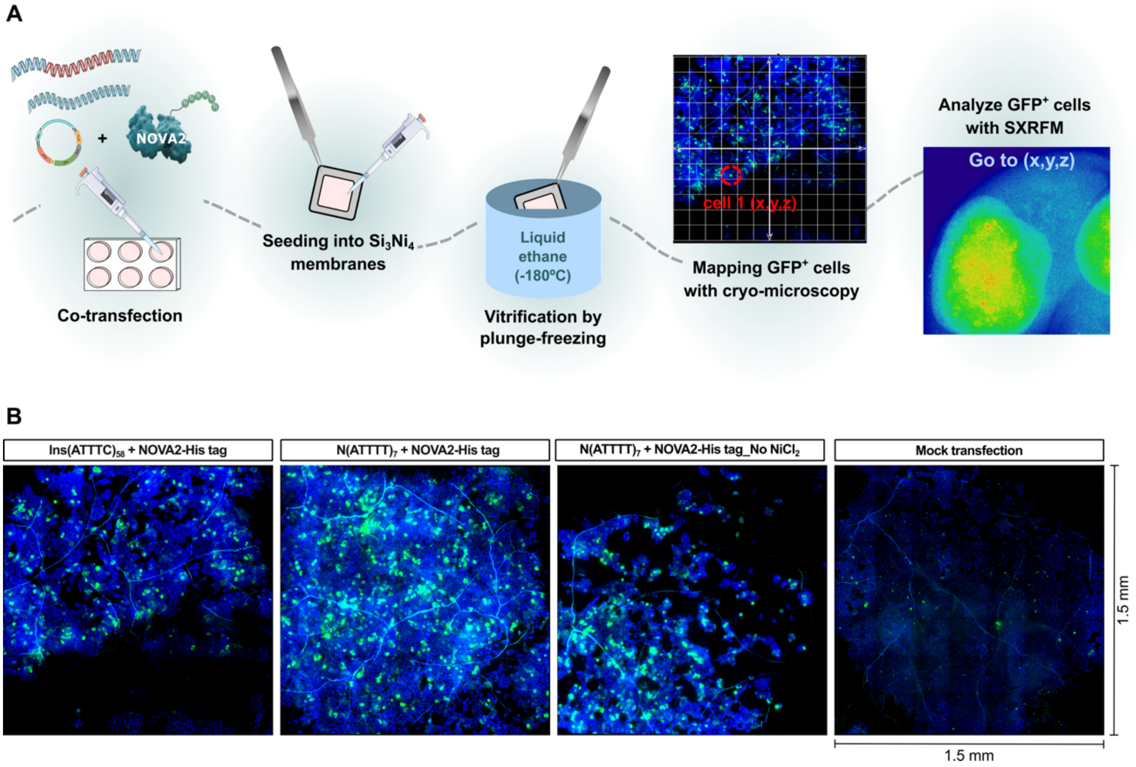


**Supplementary Figure 3 – Cryo-microscopy images of cryofixed membranes showing HEK293T cells seeded on $Si_3Ni_4$ membranes under various expression and**

**supplementation conditions.** **(A)** Schematic representation of the co-transfection, seeding into $Si_3Ni_4$ membranes, vitrification, cryo-microscopy and mapping of GFP-positive cells for SXRFM analysis. **(B)** The positions of transfected GFP-positive cells on the membranes were mapped to guide image acquisition with the high-resolution SXRFM. Images were captured using a cryogenic-fluorescence microscope (Leica EM Thunder) equipped with a HC PL APO ×50/0.9 NA cryo-objective at 88K.

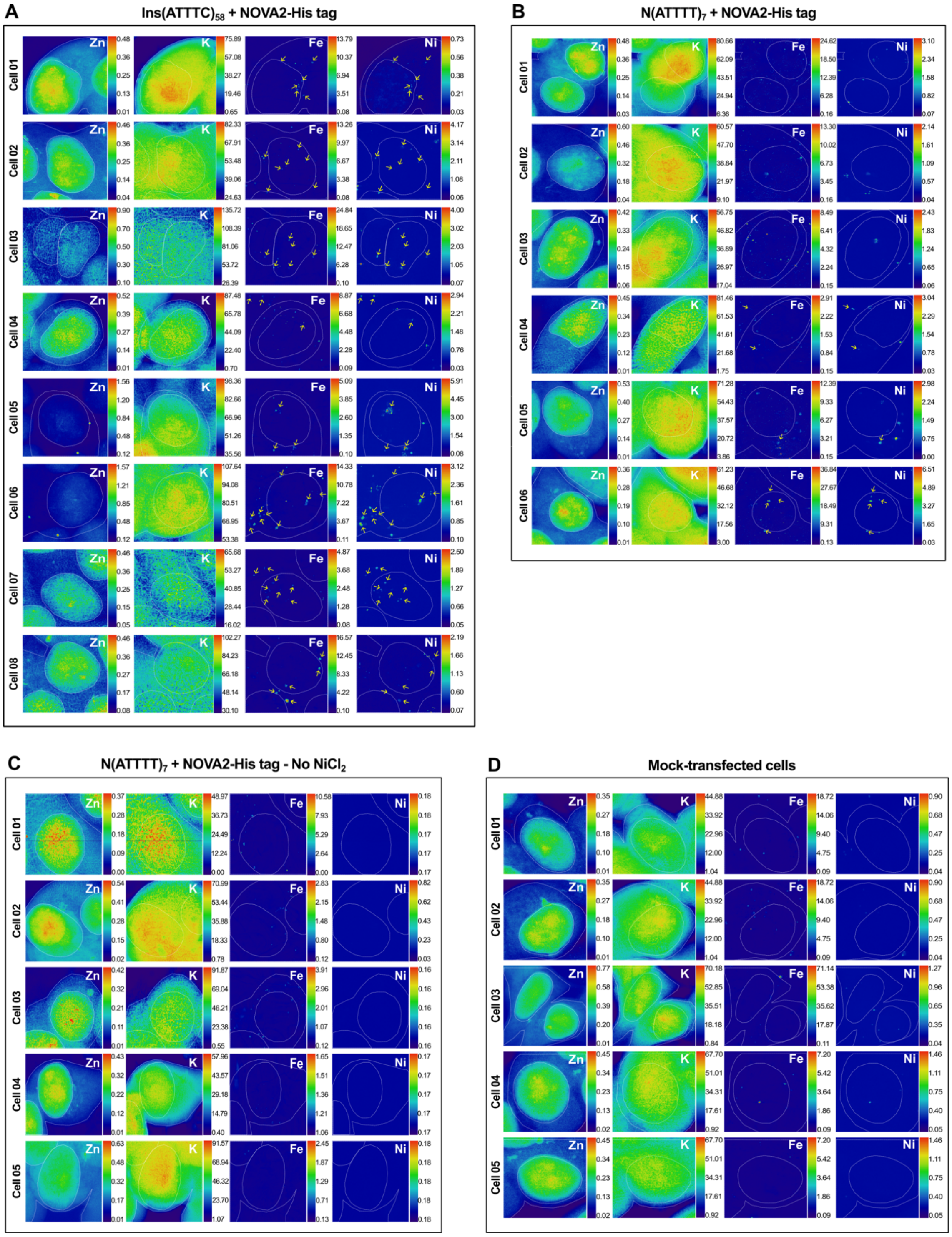


**Supplementary Figure 4 – Higher nickel-iron colocalization hotspots in Ins(ATTTC)$_{58}$-expressing cells by SXRFM.** HEK293T cells co-overexpressing NOVA2-His-tag and **(A)** the pathogenic Ins(ATTTC)$_{58}$; **(B)** the non-pathogenic N(ATTTT)$_7$; **(C)** non-pathogenic N(ATTTT)$_7$, without $NiCl_2$ supplementation and **(D)**

mock-transfected cells (no plasmids). The fluorescence images highlight the localization and expression patterns of iron (Fe) and nickel (Ni) under each condition. Zinc (Zn) was used to delineate the nucleus and potassium (K) the cytoplasm. These high-resolution nanoscopic analyses were acquired with 50 nm × 50 nm and 50 $ms.pixel^{-1}$ dwell time. The X-Ray beam size was approximately 35 nm (horizontal) x 38 nm (vertical) at an excitation energy of 17.1 keV, with an X-Ray flux of $2x10^{11}$ $photons.s^{-1}$.

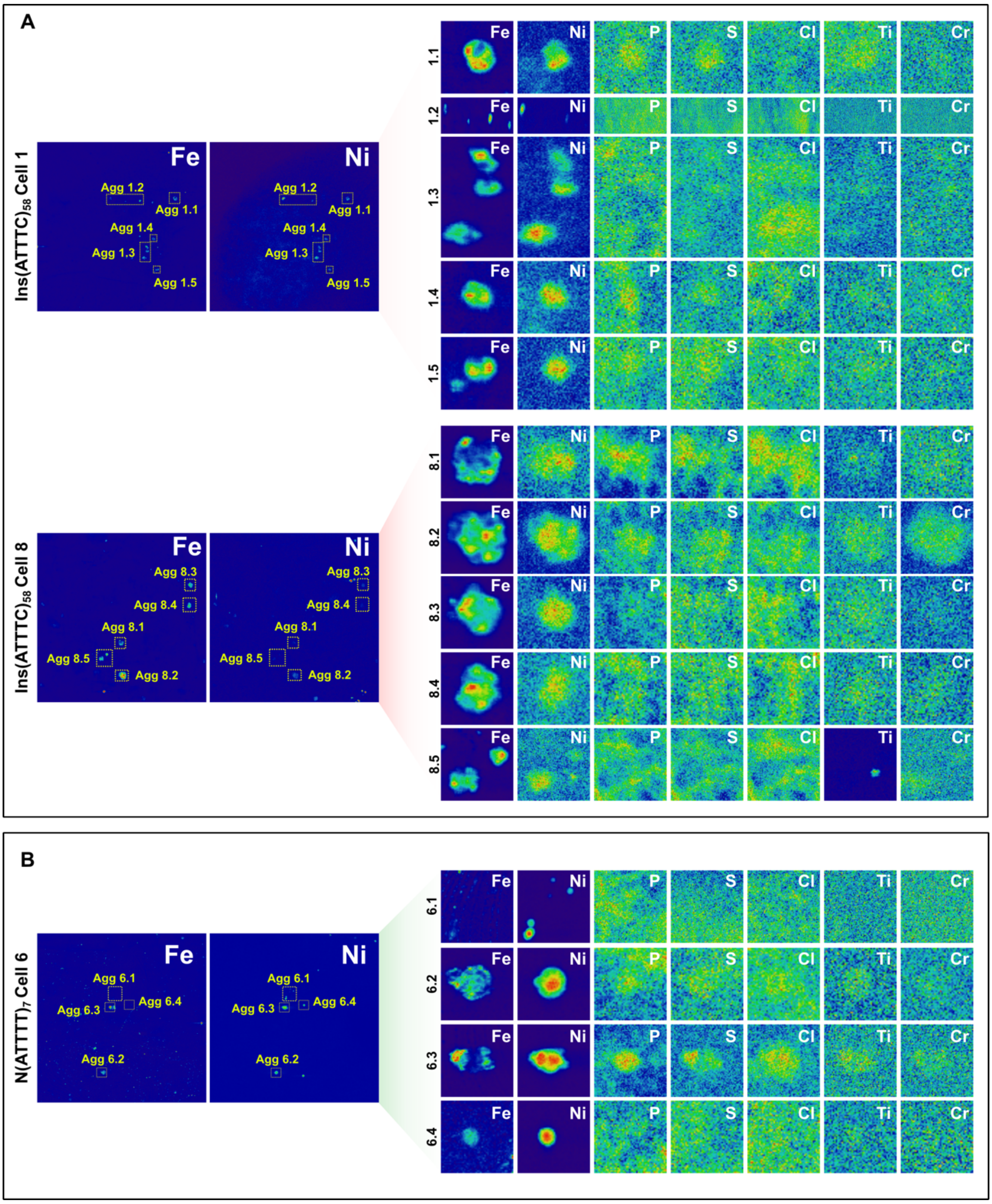


**Supplementary Figure 5 – Super high-resolution of synchrotron X-Ray fluorescence showing the different morphologies of the iron-nickel aggregates.** Regions of interest, specifically iron- and nickel-rich hotspots, were scanned with even finer resolution (20 nm × 20 nm, and 50 ms.pixel$^{-1}$ dwell time) for **(A)** Ins(ATTTC)$_{58}$-expressing cells and **(B)** N(ATTTT)$_7$-expressing cells. The X-Ray beam size was approximately 35 nm (horizontal) × 38 nm (vertical) at an excitation energy of 17.1 keV, with an X-Ray flux of $2x10^{11}$ photons.$s^{-1}$.

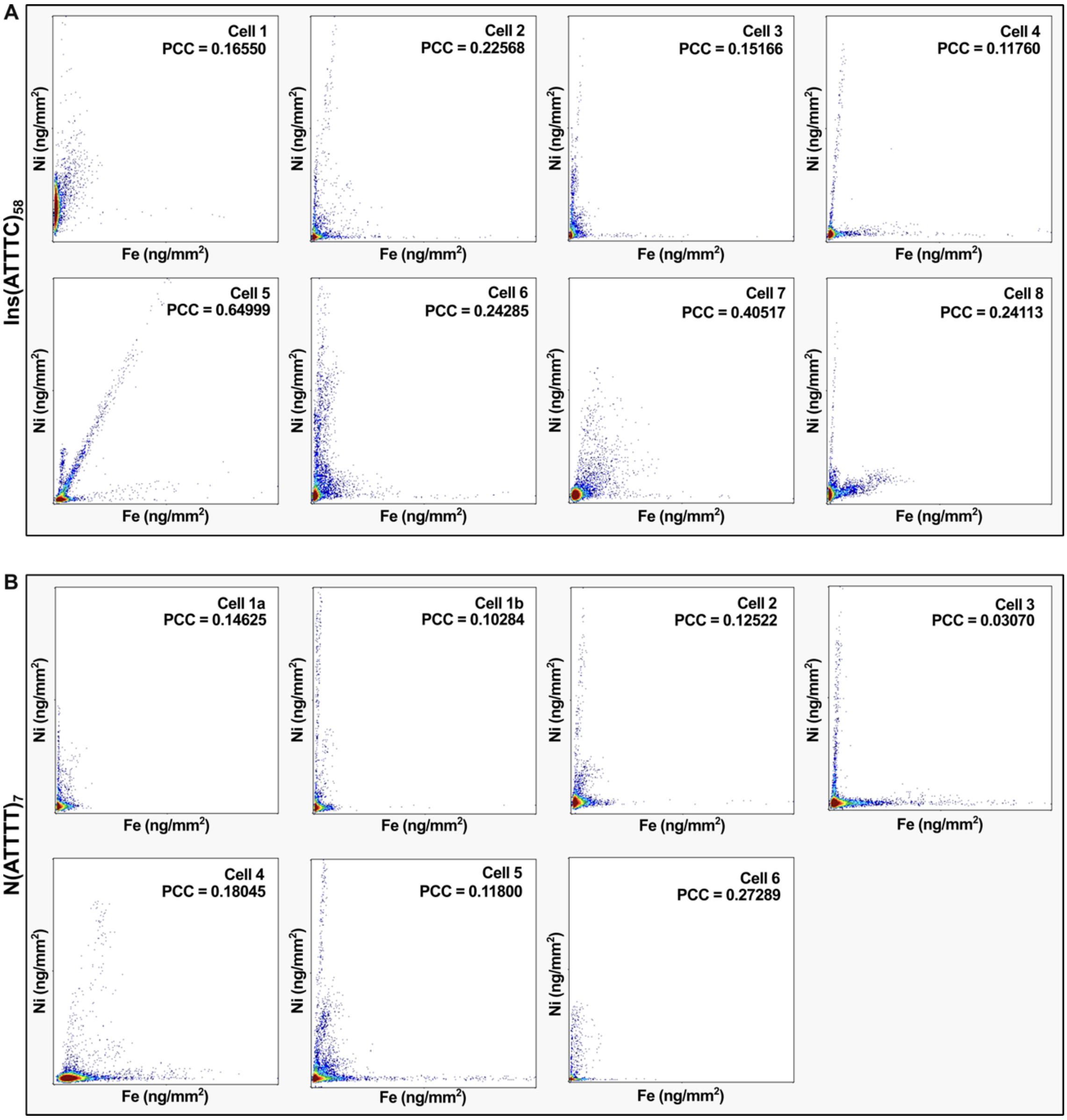


**Supplementary Figure 6 – Scatterplots illustrating the relationship between nickel (Ni) and iron (Fe) concentrations (ng/mm²) in individual cells under different conditions. (A)** Cells transfected with the Ins(ATTTC)$_{58}$ repeat and the NOVA2-His tag vector. **(B)** Cells transfected with the N(ATTTT)$_7$ repeat and the NOVA2-His tag vector. The Pearson's Correlation Coefficient (PCC) is displayed in each scatterplot, reflecting the correlation across all analyzed cells. Scatterplots and PCC calculations were generated using the ScatterJ plugin in ImageJ.

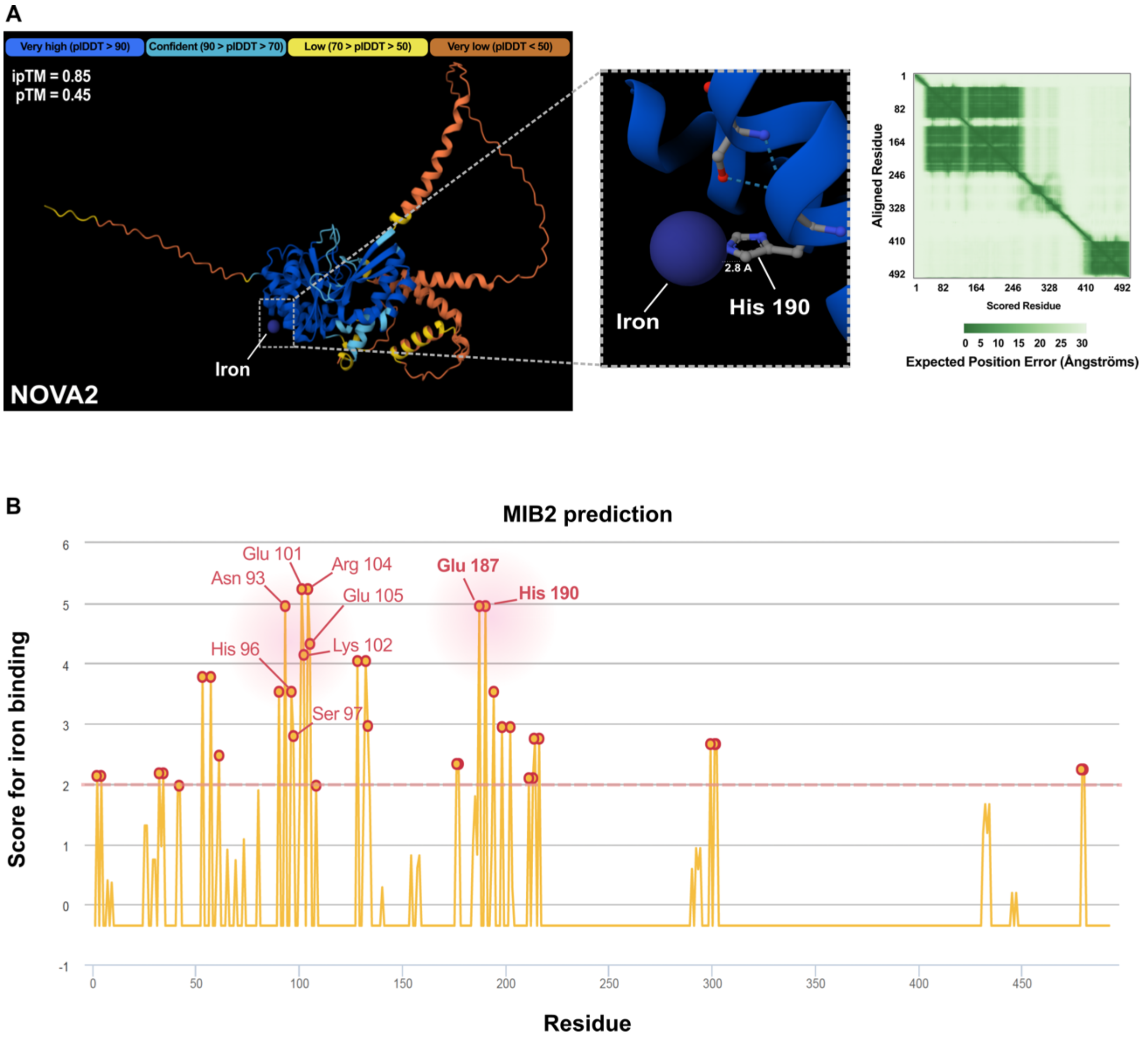


**Supplementary Figure 7 – Iron is predicted to bind NOVA2. (A)** Structural prediction of iron binding to NOVA2 by the AlphaFold 3 server. The iron ion ($Fe^{3+}$) is depicted as a blue sphere. Model confidence is color-coded: dark blue indicates pIDDT > 90, light blue for 90 > pIDDT > 70, yellow for 70 > pIDDT > 50, and orange for pIDDT < 50. The ipTM and pTM values are represented in the image and the graphical representation of the expected positional error in the AlphaFold 3 prediction is also shown on the right. **(B)** Prediction of iron-binding residues in NOVA2 with the MIB2 predictor. Red dots highlight residues with an iron-binding score > 2. Labelled residue (His190), with score exceeding 2 as determined by the MIB2 predictor, corresponds to the prediction by AlphaFold 3.

**Supplementary Table 1** - RNA-binding proteins with a higher abundance ratio in the AUUUC repeat RNA.

| Accession Number | Protein Name | Abundance Ratio (AUUUC/AUUUU) Adjusted p-value |
|---|---|---|
| E5KNY5* | Leucine-rich PPR-motif containing | 2.15e-04 |
| P19338* | Nucleolin | 1.76e-07 |
| A8MXP9* | Matrin-3 | 7.55e-12 |
| O00425 | IGF2BP3 | 3.59e-16 |
| O43390* | HNRNPR | 8.60e-12 |
| P26599* | PTBP1 | 1.49e-12 |
| Q53F64* | hnRNP AB isoform a | 2.56e-03 |
| A1A4E9 | Keratin 13 | 1.08e-08 |
| Q9NZB2 | FAM120A | 3.25e-07 |
| P19013 | Keratin 4 | 2.00e-05 |
| A0A0S2Z6K1* | HNRPLL isoform 1 | 3.59e-16 |
| B2R959 | HNRPL-like | 3.59e-16 |
| Q9UNW9* | NOVA2 | 3.59e-16 |
| P38159 | RBMX | 3.63e-12 |
| O95758* | PTBP3 | 4.64e-12 |
| Q9GZT3* | SLIRP | 5.57e-07 |
| A0A0A6YYL6 | RPL17-C18orf32 | 7.15e-04 |
| P51513* | NOVA1 | 9.69e-12 |
| Q07021 | C1QBP | 9.91e-06 |
| P08779 | Keratin 16 | 1.03e-13 |
| Q01081 | U2AF1 | 1.54e-04 |
| Q96KR1* | ZFR | 2.46e-08 |
| I3L504 | EIF5A | 2.98e-06 |
| B5BU38 | ANXA1 | 3.23e-09 |
| A8K4W5 | ACAT2-like | 9.95e-09 |
| Q8ND56 | LSM14A | 6.30e-03 |
| B4E1U9 | CDC42-like | 9.88e-06 |
| P61626 | Lysozyme C | 1.58e-05 |

*RBPs represented in Figure 3A.
